# Numerical modeling of senile plaque development under conditions of limited diffusivity of amyloid-β monomers

**DOI:** 10.1101/2024.01.09.574935

**Authors:** Andrey V. Kuznetsov

## Abstract

This paper introduces a method to simulate the progression of senile plaques, focusing on scenarios where concentrations of amyloid beta (Aβ) monomers and aggregates vary between neurons. Extracellular variations in these concentrations may arise due to limited diffusivity of Aβ monomers and a high rate of Aβ monomer production at lipid membranes, requiring a substantial concentration gradient for diffusion-driven transport of Aβ monomers. The dimensionless formulation of the model is presented, identifying four key dimensionless parameters governing the solutions for Aβ monomer and aggregate concentrations, as well as the radius of a growing Aβ plaque within the control volume. These parameters include the dimensionless diffusivity of Aβ monomers, the dimensionless rate of Aβ monomer production, and the dimensionless half-lives of Aβ monomers and aggregates. A dimensionless parameter is introduced to assess the validity of the lumped capacitance approximation. An approximate solution is derived for the scenario involving large diffusivity of Aβ monomers and dysfunctional protein degradation machinery, resulting in infinitely long half-lives for Aβ monomers and aggregates. In this scenario, the concentrations of Aβ aggregates and the radius of the Aβ plaque depend solely on a single dimensionless parameter that characterizes the rate of Aβ monomer production. According to the approximate solution, the concentration of Aβ aggregates is linearly dependent on the rate of monomer production, and the radius of an Aβ plaque is directly proportional to the cube root of the rate of monomer production. However, when departing from the conditions of the approximate solution (e.g., finite half-lives), the concentrations of Aβ monomers and aggregates, along with the plaque radius, exhibit complex dependencies on all four dimensionless parameters. For instance, under physiological half-life conditions, the plaque radius reaches a maximum value and stabilizes thereafter.

## 1. Introduction

Modeling the progression of amyloid-beta (Aβ) plaques poses a significant challenge in the field of mathematical biology, driven by their crucial role as a hallmark of Alzheimer’s disease (AD). AD is a severe neurodegenerative disorder that leads to dementia, impacting over 30 million individuals globally (Hardy, 2006; Maqbool et al., 2016; Hung and Fu, 2017; Nalivaeva and Turner, 2013). Characterized by the presence of Aβ plaques and neurofibrillary tangles containing the tau protein (Murphy and LeVine, 2010; Knopman et al., 2021), AD’s exact relationship with Aβ plaques remains uncertain. These plaques might act as causative agents, benign markers, or even protective factors in the disease (Hardy and Selkoe, 2002; O’Brien and Wong, 2011; Selkoe and Hardy, 2016; Huang et al., 2021). Rischel et al. (2023) suggested that the accumulation of Aβ serves as a protective mechanism that eventually fails.

Despite this ambiguity, the gradual accumulation of Aβ plaques precedes irreversible neurodegeneration (Yada and Naoki, 2023). A comprehensive understanding of the temporal dynamics of Aβ plaque growth is crucial for the early diagnosis and treatment of AD (Jagust et al., 2021).

The formation of an Aβ plaque is a complex process that encompasses the production of Aβ monomers, their diffusion-driven transport within the cytosol, transformation into Aβ aggregates through nucleation and autocatalytic mechanisms, proteolytic degradation of monomers and aggregates, and the assembly of Aβ aggregates into a cohesive Aβ plaque.

Aβ monomers result from the cleavage of amyloid precursor protein (APP) by β- and γ-secretases (O’Brien and Wong, 2011; Chen et al., 2017; Hampel et al., 2021). Following their release into the extracellular environment, Aβ monomers undergo diffusion in the cytosol (Waters, 2010; Bora and Prabhakara, 2009). Extracellular Aβ monomers possess the potential to aggregate (Rahman and Lendel, 2021), which was simulated in Kuznetsov (2024) using the Finke-Watzky (F-W) model (Morris et al., 2008; Iashchishyn et al., 2017). It was assumed that all Aβ aggregates become part of Aβ plaques, and the process of plaque formation from aggregates was considered sufficiently fast, making it a non-limiting stage in the model. The proteolytic degradation of Aβ monomers and aggregates was simulated by assuming finite half-lives for these components.

This paper aims to build upon previous investigations into the effect of Aβ monomer diffusivity Kuznetsov (2023) and studies based on the lumped capacitance approximation of Aβ plaque growth Kuznetsov (2024). The objective is to develop a method for predicting the growth of Aβ plaques in scenarios where the concentrations of Aβ monomers and aggregates vary across the control volume (CV), thereby not adhering to the lumped capacitance assumption. The model enables the analysis of factors such as Aβ monomer production, diffusion-driven transport of Aβ monomers, the kinetic rate of conversion of Aβ monomers into aggregates, and the half-lives of Aβ monomers and aggregates due to proteolytic degradation. The model is presented in a dimensionless form to minimize the number of parameters influencing its solution. The impact of each dimensionless parameter on the concentrations of Aβ monomers and aggregates, as well as the radius of the Aβ plaque, is examined.

## 2. Materials and models

### 2.1. Model equations

The F-W model is employed to simulate the conversion of Aβ monomers into aggregates. This conversion takes place through two mechanisms: nucleation and autocatalysis. During the nucleation step, new aggregates are continuously formed (as described in Eq. (1) below), and in the autocatalysis step, there is rapid surface growth of these aggregates (as described in Eq. (2) below) (Morris et al., 2008; Iashchishyn et al., 2017):

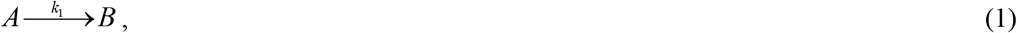

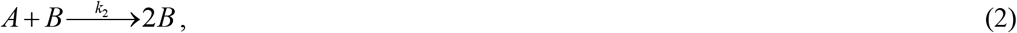

where *A* represents a monomeric protein, and *B* represents an amyloid-converted protein. The kinetic constants, *k*_1_ and *k*_2_, represent the rates of nucleation and autocatalytic growth, respectively (Morris et al., 2008). Eqs. (1) and (2) describe pseudo-elementary reactions, which, in reality, involve a much larger number of reaction steps.

The typical application of the F-W model occurs when a given initial concentration of monomers is converted into aggregates (Iashchishyn et al., 2017). In this paper, the model is applied to a situation where monomers are continuously supplied to the CV through a boundary located at *x*=0 (see Fig. 1). The generation of monomers occurs through the cleavage of APP by β- and γ-secretases at lipid membranes, such as a cellular membrane (Haass et al., 2012; Masters and Selkoe, 2012; O’Brien and Wong, 2011; Hampel et al., 2021).

**Fig. 1.**
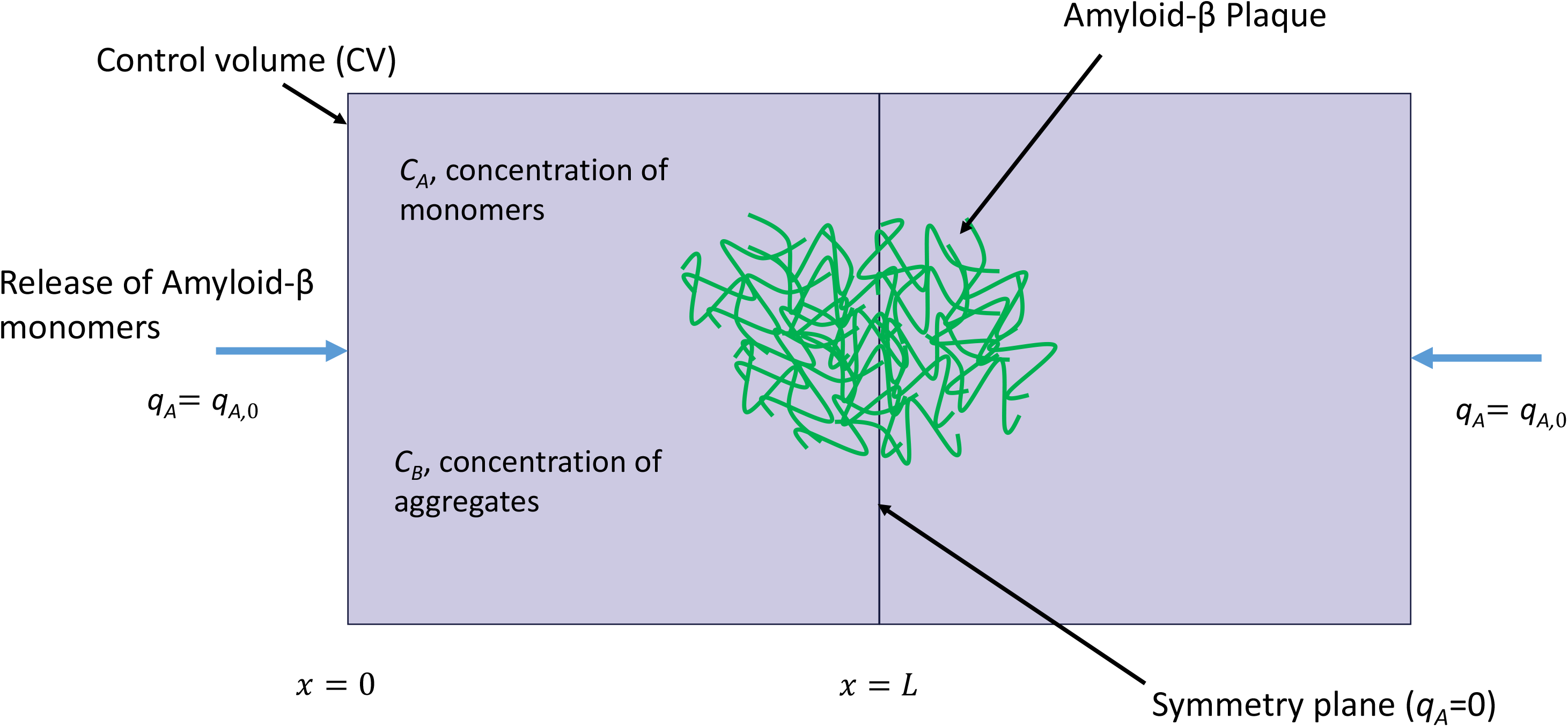
A schematic diagram illustrating two CVs. An Aβ plaque forms at the interface between the CVs. Aβ monomers enter the CV from one of its faces, and Aβ aggregates are produced through a combined process of nucleation and autocatalytic growth. Aβ monomers are generated at a lipid membrane (assumed at the *x*=0 boundary). The *x*=*L* boundary is modeled as symmetric, resulting in no monomer flux through it.

Expressing the conservation of Aβ monomers yields:

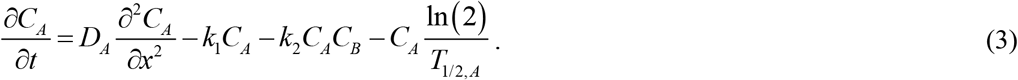

Expressing the conservation of Aβ aggregates yields:

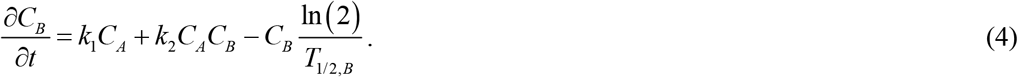

The initial conditions are described by the following equations:

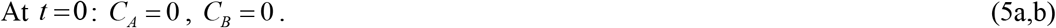

The boundary conditions are expressed through the following equations:

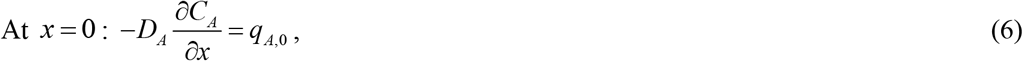

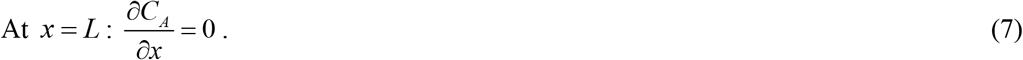

Eq. (6) posits that Aβ monomers are released from the boundary located at *x*=0. The boundary at *x*=*L* is modeled as a symmetric one. The model employs the independent variables listed in Table 1, while Table 2 outlines the dependent variables. Table 3 gives a summary of the parameters used in the model.

**Table 1.**
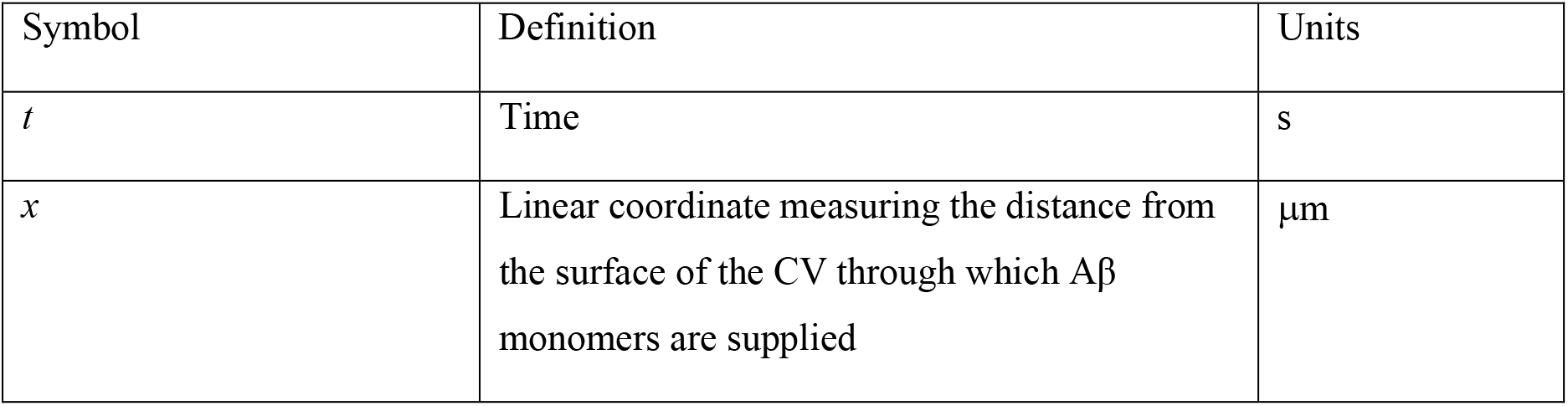
Independent variables employed in the model.

| Symbol | Definition | Units |
| --- | --- | --- |
| $t$ | Time | s |
| $x$ | Linear coordinate measuring the distance from the surface of the CV through which Aβ monomers are supplied | $\mu\text{m}$ |

**Table 2.**
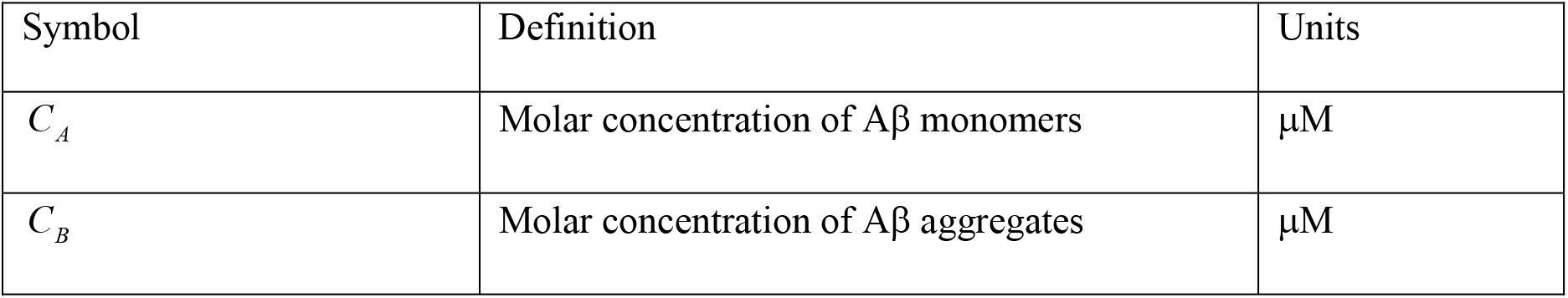
Dependent variables used in the model.

| Symbol | Definition | Units |
| --- | --- | --- |
| $C_A$ | Molar concentration of Aβ monomers | $\mu\text{M}$ |
| $C_B$ | Molar concentration of Aβ aggregates | $\mu\text{M}$ |

**Table 3.**
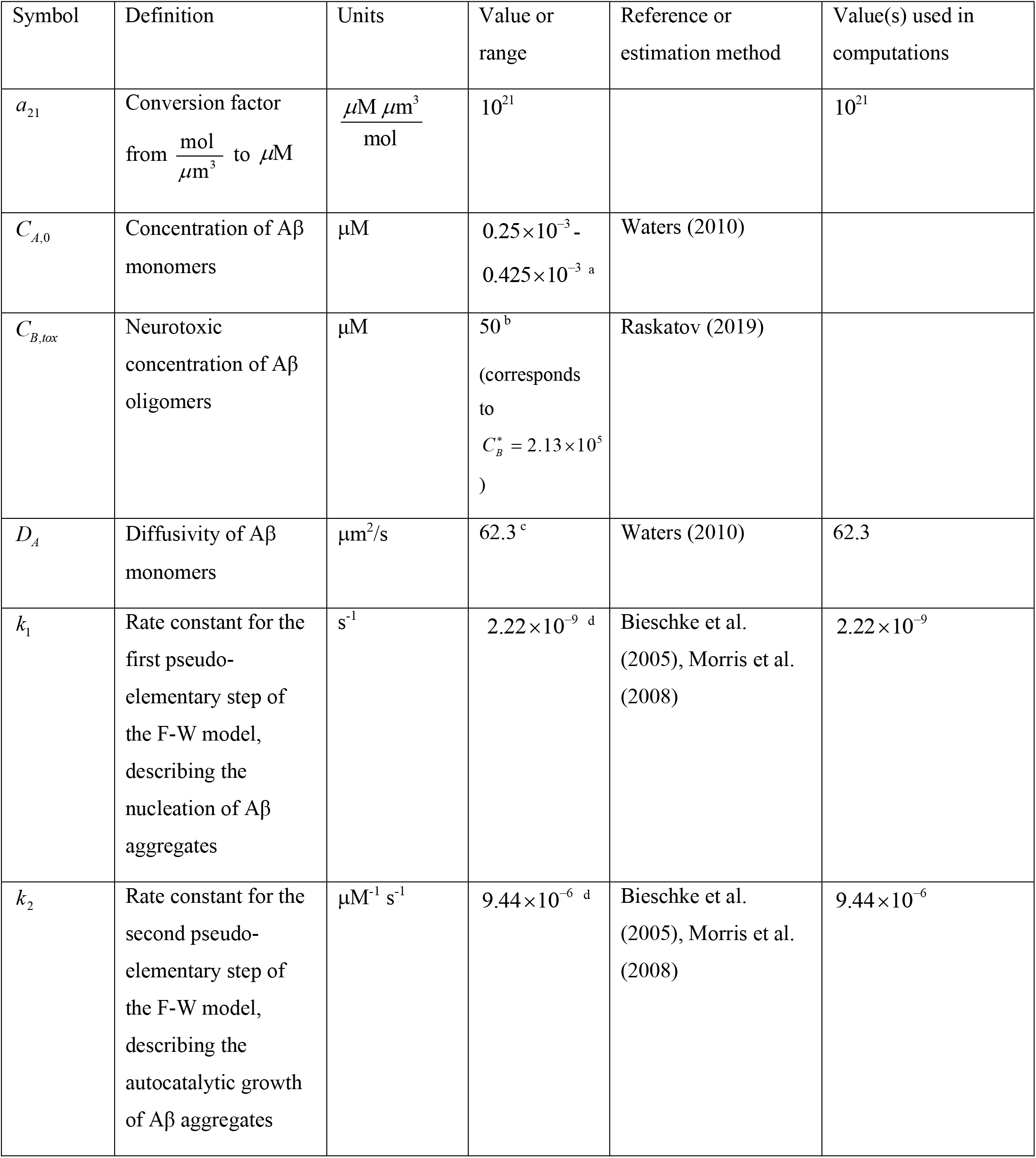
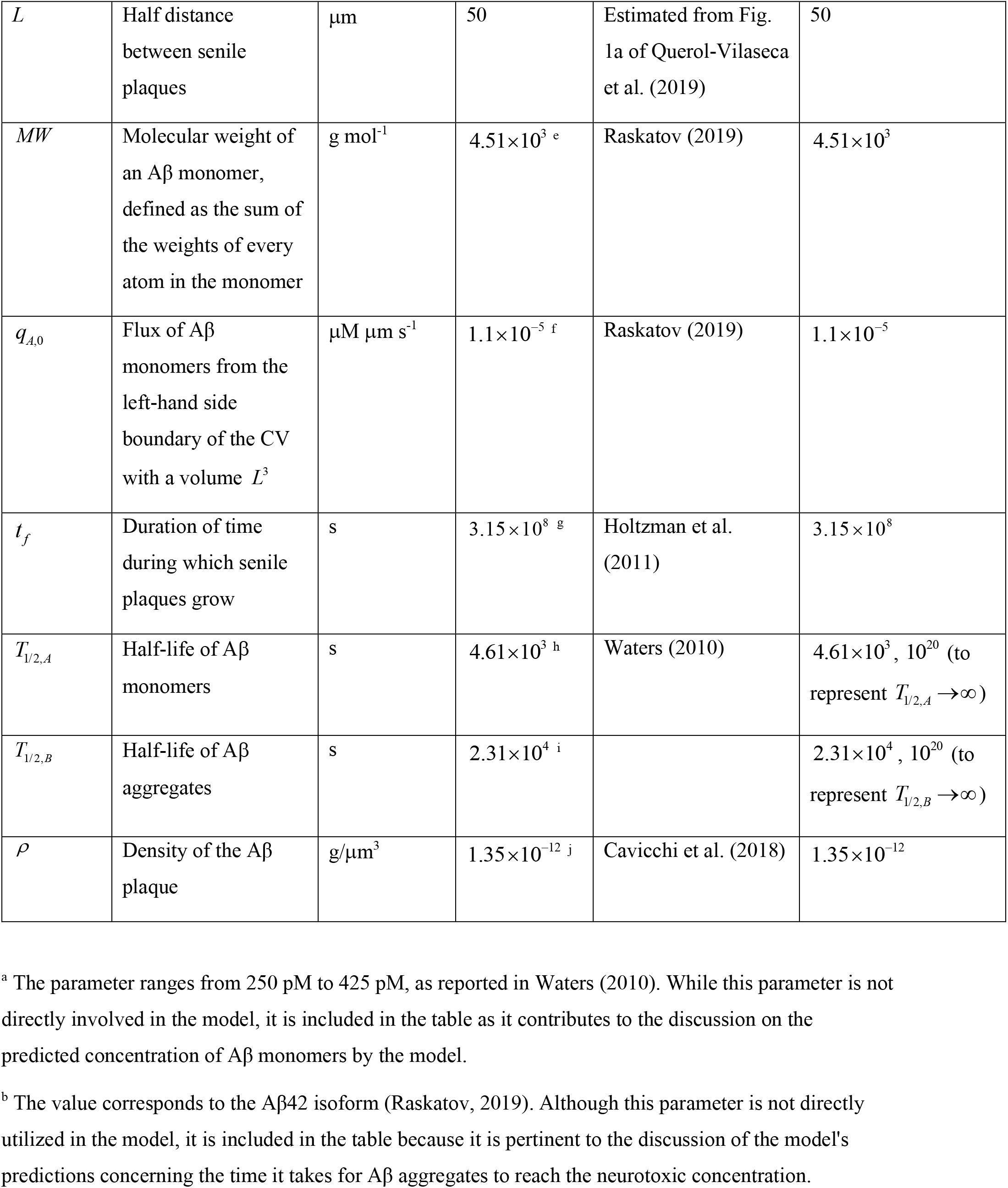
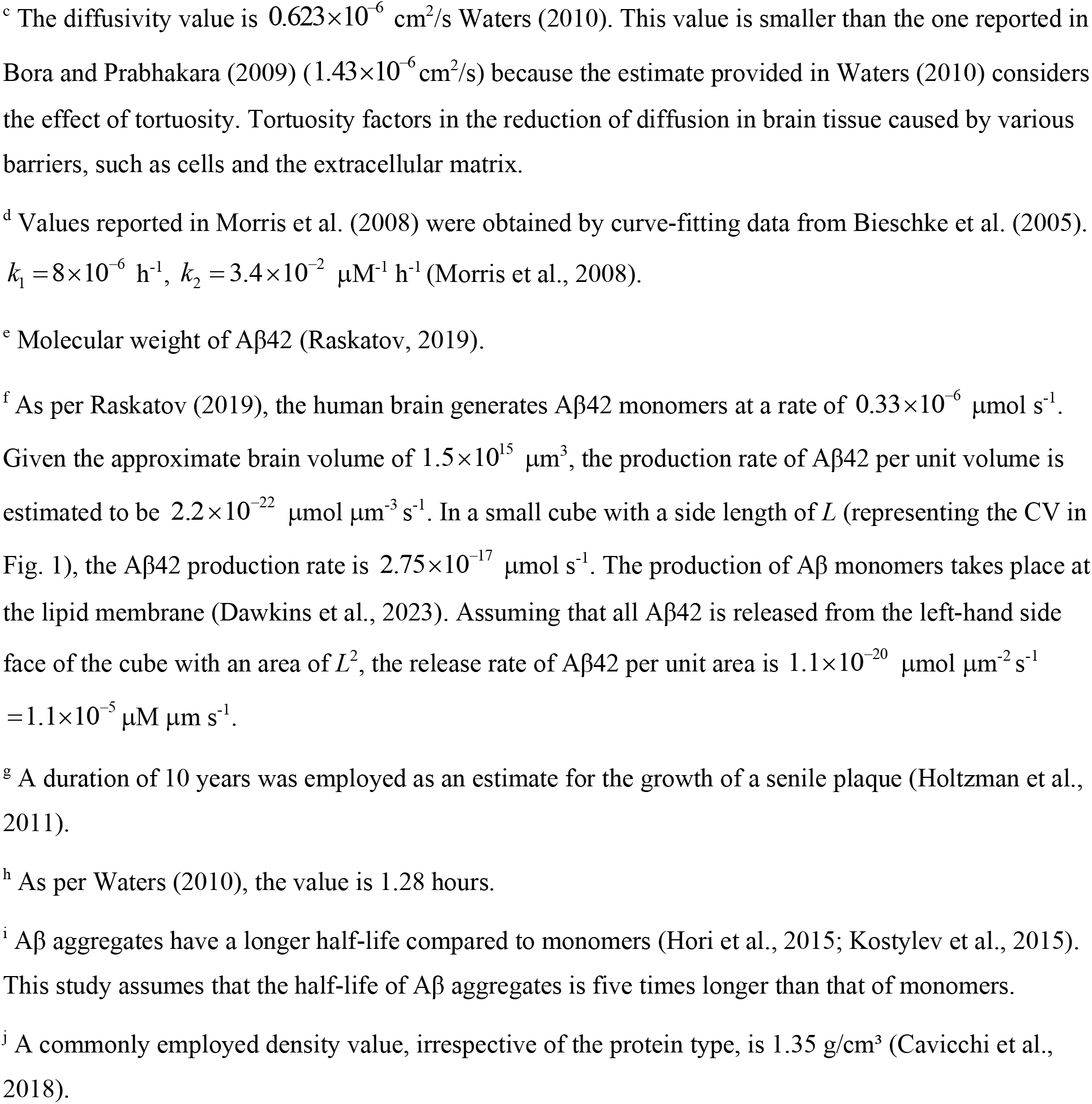
Parameters utilized in the model.

| Symbol | Definition | Units | Value or range | Reference or estimation method | Value(s) used in computations |
| --- | --- | --- | --- | --- | --- |
| $a_{21}$ | Conversion factor from $\frac{\text{mol}}{\mu\text{m}^3}$ to $\mu\text{M}$ | $\frac{\mu\text{M } \mu\text{m}^3}{\text{mol}}$ | $10^{21}$ | | $10^{21}$ |
| $C_{A,0}$ | Concentration of Aβ monomers | $\mu\text{M}$ | $0.25 \times 10^{-3}$ - $0.425 \times 10^{-3}$ <sup>a</sup> | Waters (2010) | |
| $C_{B,tox}$ | Neurotoxic concentration of Aβ oligomers | $\mu\text{M}$ | $50$ <sup>b</sup><br>(corresponds to $C_B^* = 2.13 \times 10^5$) | Raskatov (2019) | |
| $D_A$ | Diffusivity of Aβ monomers | $\mu\text{m}^2/\text{s}$ | $62.3$ <sup>c</sup> | Waters (2010) | 62.3 |
| $k_1$ | Rate constant for the first pseudo-elementary step of the F-W model, describing the nucleation of Aβ aggregates | $\text{s}^{-1}$ | $2.22 \times 10^{-9}$ <sup>d</sup> | Bieschke et al. (2005), Morris et al. (2008) | $2.22 \times 10^{-9}$ |
| $k_2$ | Rate constant for the second pseudo-elementary step of the F-W model, describing the autocatalytic growth of Aβ aggregates | $\mu\text{M}^{-1} \text{s}^{-1}$ | $9.44 \times 10^{-6}$ <sup>d</sup> | Bieschke et al. (2005), Morris et al. (2008) | $9.44 \times 10^{-6}$ |
| $L$ | Half distance between senile plaques | $\mu\text{m}$ | 50 | Estimated from Fig. 1a of Querol-Vilaseca et al. (2019) | 50 |
| $MW$ | Molecular weight of an Aβ monomer, defined as the sum of the weights of every atom in the monomer | $\text{g mol}^{-1}$ | $4.51 \times 10^3$ <sup>e</sup> | Raskatov (2019) | $4.51 \times 10^3$ |
| $q_{A,0}$ | Flux of Aβ monomers from the left-hand side boundary of the CV with a volume $L^3$ | $\mu\text{M } \mu\text{m s}^{-1}$ | $1.1 \times 10^{-5}$ <sup>f</sup> | Raskatov (2019) | $1.1 \times 10^{-5}$ |
| $t_f$ | Duration of time during which senile plaques grow | s | $3.15 \times 10^8$ <sup>g</sup> | Holtzman et al. (2011) | $3.15 \times 10^8$ |
| $T_{1/2,A}$ | Half-life of Aβ monomers | s | $4.61 \times 10^3$ <sup>h</sup> | Waters (2010) | $4.61 \times 10^3$, $10^{20}$ (to represent $T_{1/2,A} \rightarrow \infty$) |
| $T_{1/2,B}$ | Half-life of Aβ aggregates | s | $2.31 \times 10^4$ <sup>i</sup> | | $2.31 \times 10^4$, $10^{20}$ (to represent $T_{1/2,B} \rightarrow \infty$) |
| $\rho$ | Density of the Aβ plaque | $\text{g}/\mu\text{m}^3$ | $1.35 \times 10^{-12}$ <sup>j</sup> | Cavicchi et al. (2018) | $1.35 \times 10^{-12}$ |

<sup>a</sup> The parameter ranges from 250 pM to 425 pM, as reported in Waters (2010). While this parameter is not directly involved in the model, it is included in the table as it contributes to the discussion on the predicted concentration of Aβ monomers by the model.
<sup>b</sup> The value corresponds to the Aβ 42 isoform (Raskatov, 2019). Although this parameter is not directly utilized in the model, it is included in the table because it is pertinent to the discussion of the model's predictions concerning the time it takes for Aβ aggregates to reach the neurotoxic concentration.
<sup>c</sup> The diffusivity value is $0.623 \times 10^{-6} \text{ cm}^2/\text{s}$ Waters (2010). This value is smaller than the one reported in Bora and Prabhakara (2009) ( $1.43 \times 10^{-6} \text{ cm}^2/\text{s}$ ) because the estimate provided in Waters (2010) considers the effect of tortuosity. Tortuosity factors in the reduction of diffusion in brain tissue caused by various barriers, such as cells and the extracellular matrix.
<sup>d</sup> Values reported in Morris et al. (2008) were obtained by curve-fitting data from Bieschke et al. (2005). $k_1 = 8 \times 10^{-6} \text{ h}^{-1}$ , $k_2 = 3.4 \times 10^{-2} \text{ } \mu\text{M}^{-1} \text{ h}^{-1}$ (Morris et al., 2008).
<sup>e</sup> Molecular weight of Aβ 42 (Raskatov, 2019).
<sup>f</sup> As per Raskatov (2019), the human brain generates Aβ 42 monomers at a rate of $0.33 \times 10^{-6} \text{ } \mu\text{mol s}^{-1}$ . Given the approximate brain volume of $1.5 \times 10^{15} \text{ } \mu\text{m}^3$ , the production rate of Aβ 42 per unit volume is estimated to be $2.2 \times 10^{-22} \text{ } \mu\text{mol } \mu\text{m}^{-3} \text{ s}^{-1}$ . In a small cube with a side length of $L$ (representing the CV in Fig. 1), the Aβ 42 production rate is $2.75 \times 10^{-17} \text{ } \mu\text{mol s}^{-1}$ . The production of Aβ monomers takes place at the lipid membrane (Dawkins et al., 2023). Assuming that all Aβ 42 is released from the left-hand side face of the cube with an area of $L^2$ , the release rate of Aβ 42 per unit area is $1.1 \times 10^{-20} \text{ } \mu\text{mol } \mu\text{m}^{-2} \text{ s}^{-1} = 1.1 \times 10^{-5} \text{ } \mu\text{M } \mu\text{m s}^{-1}$ .
<sup>g</sup> A duration of 10 years was employed as an estimate for the growth of a senile plaque (Holtzman et al., 2011).
<sup>h</sup> As per Waters (2010), the value is 1.28 hours.
<sup>i</sup> Aβ aggregates have a longer half-life compared to monomers (Hori et al., 2015; Kostylev et al., 2015). This study assumes that the half-life of Aβ aggregates is five times longer than that of monomers.
<sup>j</sup> A commonly employed density value, irrespective of the protein type, is $1.35 \text{ g/cm}^3$ (Cavicchi et al., 2018).

### 2.2. A criterion characterizing the limitations of the lumped capacitance approximation

The criterion for the validity of the lumped capacitance approximation is derived for the scenario of dysfunctional degradation machinery for Aβ monomers and aggregates, i.e., *T*_1/ 2, *A*_ →∞ and *T*_1/ 2,*B*_ →∞. Utilizing this approximation and summing Eqs. (3) and (4) under steady-state conditions yields

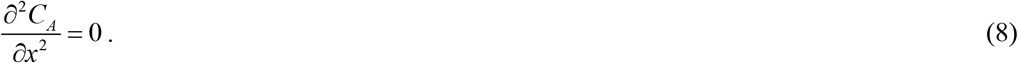

Eq. (8) is solved subject to the boundary condition given by Eq. (6) and the following additional boundary condition:

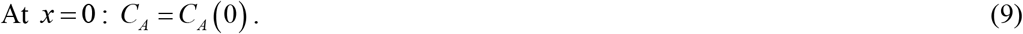

This yields the following concentration distribution of Aβ monomers:

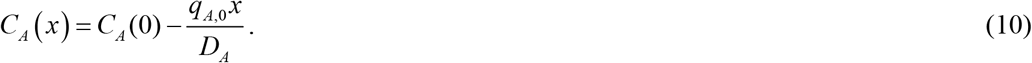

From Eq. (10), it follows that

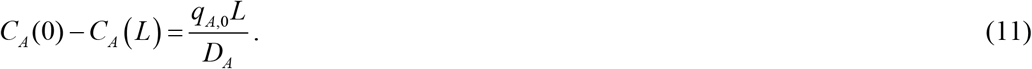

The approximate solution for *C*_*A*_ is given by Eq. (S12). This equation is derived in section S1 of the Supplemental Materials. From Eq. (S12), it follows that

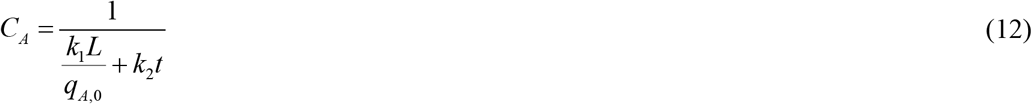

At large times, it follows from Eq. (12) that

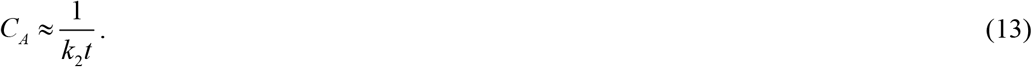

Using Eqs. (11) and (13), a parameter that characterizes the ratio of *C*_*A*_ variation across the CV to the average value of *C*_*A*_ in the CV at time *t* _*f*_ is defined as follows:

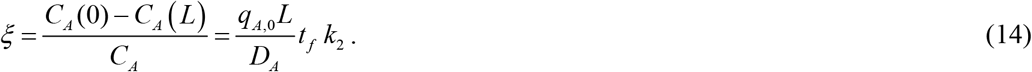

The role of parameter *ξ* is analogous to that of the Biot number in convection-conduction problems. The lumped capacitance approximation is applicable as long as *ξ* ≪ 1. It is interesting that the deviation from the lumped capacitance conditions increases as the duration of the process, *t* _*f*_, increases.

### 2.3. Dimensionless model equations

The dimensionless independent variables employed in the model are summarized in Table 4, while the dimensionless dependent variables are given in Table 5. A summary of the dimensionless parameters utilized in the model is provided in Table 6.

**Table 4.** Dimensionless independent variables employed in the model.

| Symbol | Definition |
| --- | --- |
| $t^*$ | $t k_1$ |
| $x^*$ | $x / L$ |

**Table 5.** Dimensionless dependent variables utilized in the model.

| Symbol | Definition |
| --- | --- |
| $C_A^*$ | $C_A k_2 / k_1$ |
| $C_B^*$ | $C_B k_2 / k_1$ |
| $r_{ABP}^*$ | $r_{ABP} / L$ |

**Table 6.** Dimensionless parameters utilized in the model.

| Symbol | Definition | Value(s) obtained using the dimensional parameter values listed in the final column of Table 3 |
| --- | --- | --- |
| $D_A^*$ | $D_A / (L^2 k_1)$ | $1.12 \times 10^7$ |
| $q_{A,0}^*$ | $q_{A,0} k_2 / (k_1^2 L)$ | $4.21 \times 10^5$ |
| $t_f^*$ | $t_f k_1$ | 0.700 |
| $T_{1/2,A}^*$ | $T_{1/2,A} k_1$ | $1.02 \times 10^{-5}$, $2.22 \times 10^{11}$ (to represent $T_{1/2,A}^* \rightarrow \infty$) |
| $T_{1/2,B}^*$ | $T_{1/2,B} k_1$ | $5.12 \times 10^{-5}$, $2.22 \times 10^{11}$ (to represent $T_{1/2,B}^* \rightarrow \infty$) |
| $\rho^*$ | $\frac{\rho}{MW} a_{21} \frac{k_2}{k_1}$ | $1.27 \times 10^9$ |

The model, expressed by Eqs. (3)-(7), is reformulated in its dimensionless form as follows:

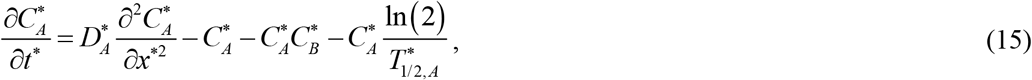

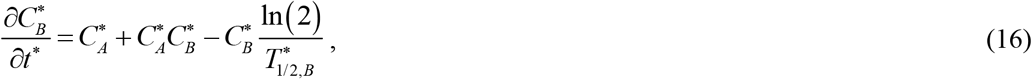

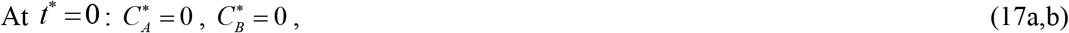

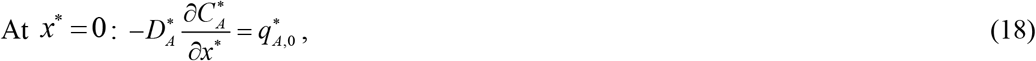

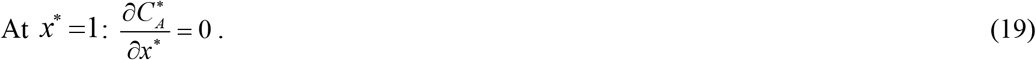

Using dimensionless parameters, parameter ξ defined in Eq. (14) can be reformulated as:

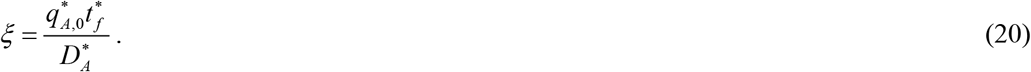

To explore the effect of different model parameters, Eqs. (15)-(19) are solved numerically. The numerical solution is then compared with the approximate solution of Eqs. (15)-(19) obtained in Kuznetsov (2024). The approximate solution, detailed in section S1 of the Supplemental Materials, is derived using the lumped capacitance approximation under the assumption of dysfunctional protein degradation machinery, implying infinitely large half-lives for Aβ monomers and aggregates (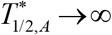 and 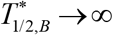).

The model developed in this paper does not simulate the process in which adhesive Aβ fibrils come together to form an Aβ plaque. It is presumed that this assembly process happens more rapidly than the creation of Aβ fibrils.

The growth of the Aβ plaque (Fig. 1) is simulated by employing the following method. The total count of Aβ monomers integrated into the Aβ plaque, represented as *N*, at a given time *t*, is calculated using the following equation (Watzky et al., 2008):

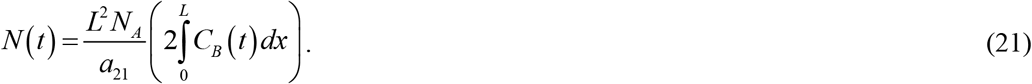

The factor of 2 preceding the integral in Eq. (21) is employed because the model posits that the Aβ plaque forms between two neurons, with each membrane releasing Aβ monomers at a rate of *q*_*A*,0_ (Fig. 1).

Alternatively, *N* (*t*) can be calculated as outlined in Watzky et al. (2008):

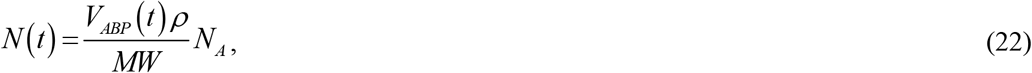

where *MW* denotes the average molecular weight of an Aβ monomer.

By setting the expressions on the right-hand sides of Eqs. (21) and (22) equal and solving for the volume of an Aβ plaque, the following result is obtained:

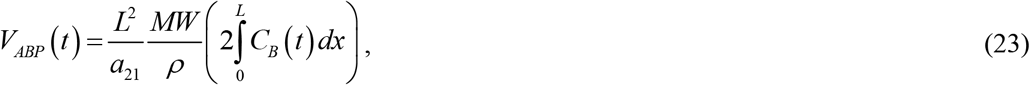

where 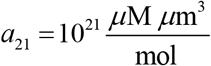 denotes the conversion factor from 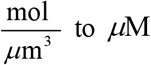.

Assuming that the Aβ plaque is spherical in shape,

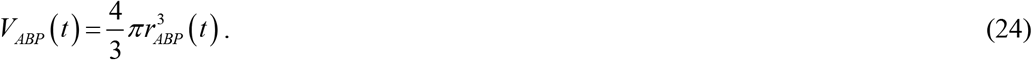

Setting the right-hand sides of Eqs. (23) and (24) equal and solving for the radius of the Aβ plaque, the following result is obtained:

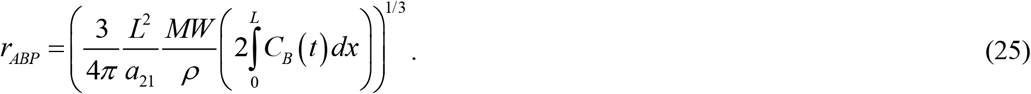

Expressed in dimensionless variables, Eq. (25) becomes:

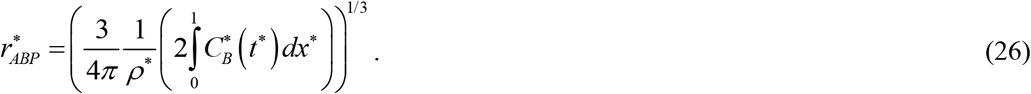

If an approximate solution for 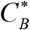 given by Eq. (S10) is used in Eq. (26), the following approximate solution for 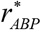 is obtained:

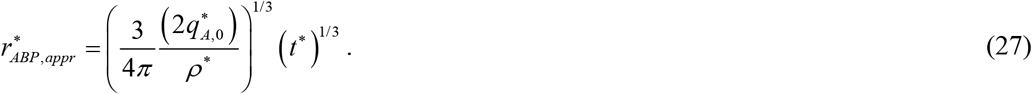

Returning to the dimensional variables, Eq. (27) can be reformulated as:

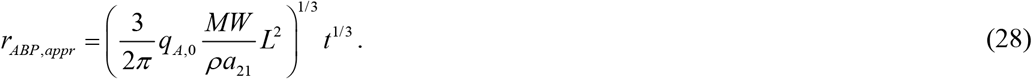

Eq. (28) suggests that the radius of an Aβ plaque is directly proportional to the cube root of the plaque’s growth time. This equation indicates that, at large times, the plaque growth is restricted by Aβ peptide production, *q*_*A*,0_, and is independent of kinetic constants, *k*_1_ and *k*_2_. This aligns with the results reported in Burgold et al. (2014) for the late stage of plaque growth, referred to in Burgold et al. (2014) as the late asymptotic phase. It should be noted that Eq. (28) (Eq. (27) in the dimensionless form) applies only to the situation when the protein degradation machinery is dysfunctional, and half-lives of Aβ monomers and aggregates are infinitely long, while Eq. (26) applies for any half-life of Aβ monomers and aggregates. In various figures, the predictions obtained from Eq. (26) are compared with predictions that follow from the approximate analytical solution (27).

Eq. (27) is valid for large times. To investigate how the radius of the Aβ plaque increases when *t*^*^ → 0, Eq. (S9) is substituted into Eq. (26). This gives the following result for the initial growth of the plaque:

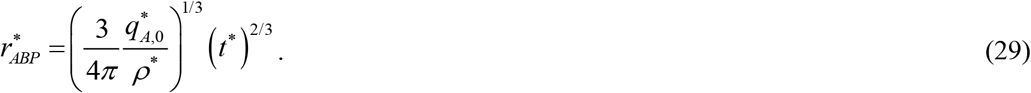

Returning to dimensional variables, Eq. (29) can be reformulated as:

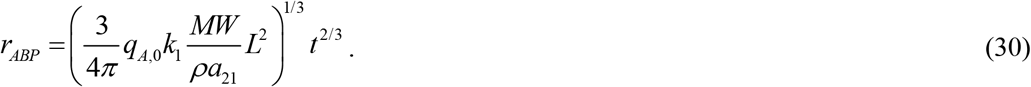

Eq. (30) indicates that during the initial growth of the plaque, its radius is influenced by the rate of nucleation of Aβ aggregates, *k*_1_.

### 2.4. Numerical solution

Eqs. (15) and (16) constitute a system of two partial differential equations (PDEs). They were solved with initial conditions given by Eq. (17) and boundary conditions given by Eqs. (18) and (19), using a well-validated MATLAB’s PDEPE solver (MATLAB R2020b, MathWorks, Natick, MA, USA). A (100,100) solution mesh was employed, except for Fig. S12, where a (1000,1000) solution mesh was used. In addition to the boundary conditions on 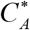, PDEPE requires boundary conditions on 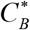 at both ends of the domain. Since Aβ aggregates are assumed to have zero diffusivity (as per Eq. (4)), zero boundary conditions on the diffusion flux of 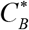 are imposed on both sides of the domain. Numerical integration of 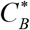 with respect to *x*^*^ in Eq. (26) was conducted using MATLAB’s TRAPZ solver, implementing the trapezoidal method.

## 3. Results

### 3.1. Dimensionless parameters

In this study, the effects of four dimensionless parameters were explored: (1) dimensionless diffusivity of Aβ monomers, 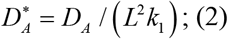 dimensionless rate at which Aβ monomers were produced, 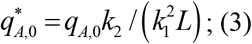 dimensionless half-life of Aβ monomers, 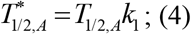 dimensionless half-life of Aβ aggregates, 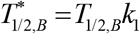. The other two parameters from Table 6 were kept constant at the following values: 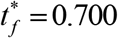 (equivalent to *t* =10 years) and *ρ* ^*^ = 1.27 ×10^9^ (equivalent to 1.35 g/cm^3^).

### 3.2. Effects of variating the dimensionless diffusivity

Effects of varying the dimensionless diffusivity of Aβ monomers, 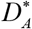, are analyzed in Figs. 2, 3, and Figs. S1-S3 in the Supplemental Materials.

**Fig. 2.**
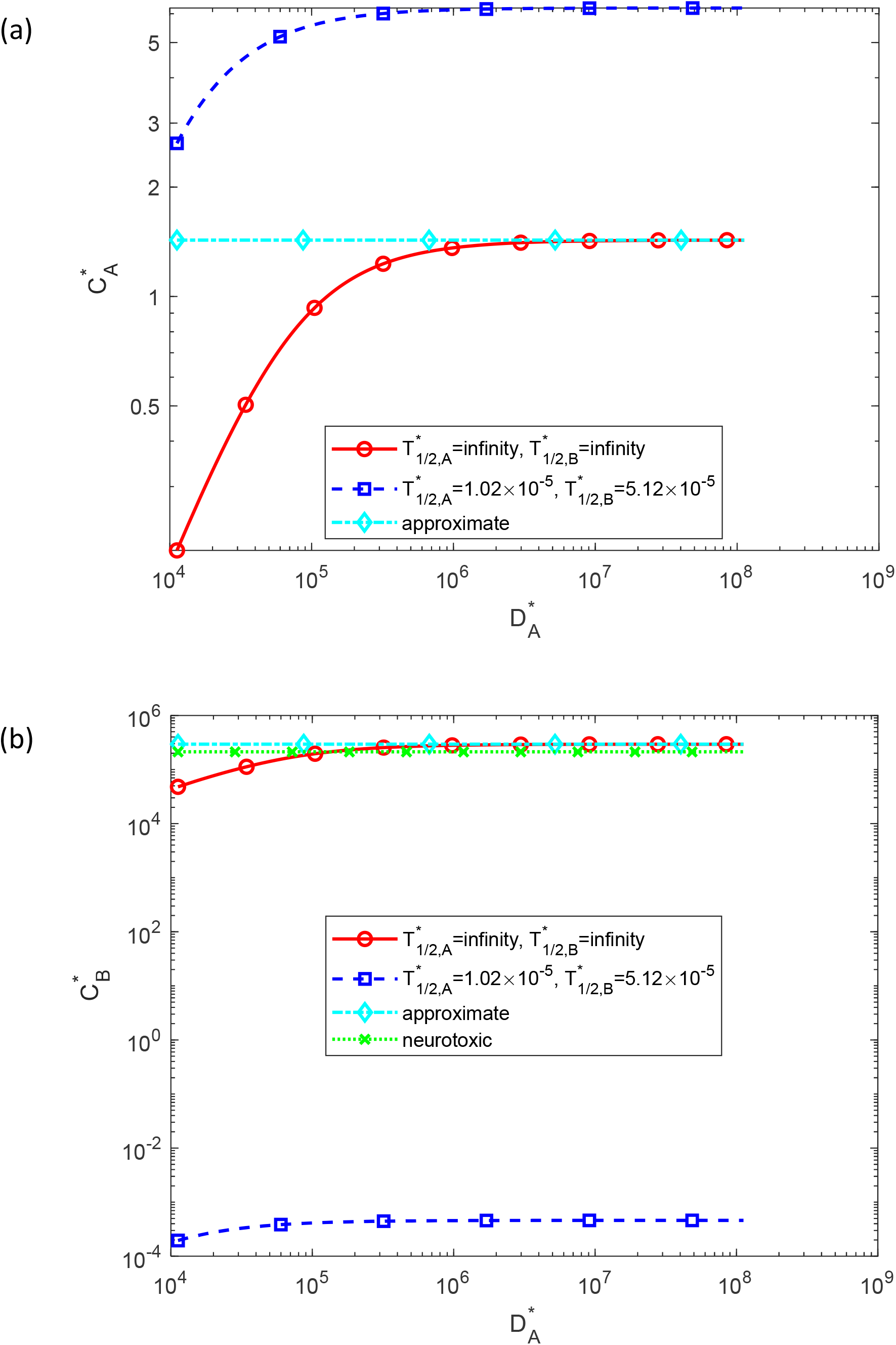
Dimensionless molar concentration of Aβ monomers, 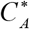 (a), and Aβ aggregates, 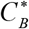 (b), plotted against the dimensionless diffusivity of Aβ monomers, 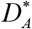. Approximate solutions for 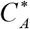 and 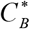 are computed using Eqs. (S11) and (S10), respectively. The computational results are presented at the right-hand side boundary of the CV, *x*^*^ =1, at 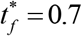 (10 years).

**Fig. 3.**
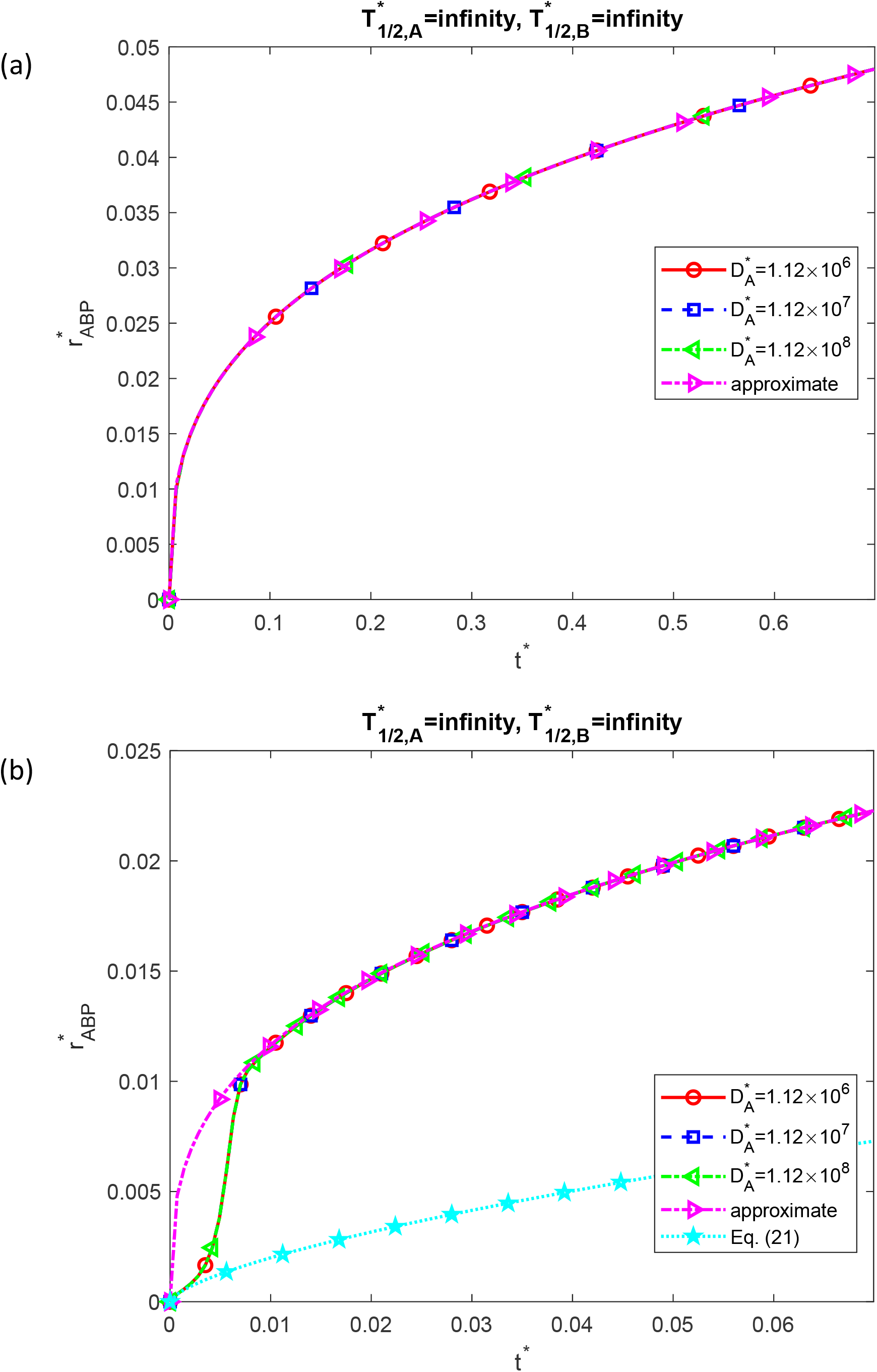
(a) Dimensionless radius of a growing Aβ plaque, 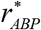, as a function of dimensionless time, *t*^*^. (b) Similar to Fig. 3a but focusing on the time range [0 0.07] on the *x*-axis. The approximate solution for 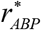 is plotted using Eq. (27); 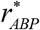 is directly proportional to (*t*^*^)^1/3^. These computational results are presented at the right-hand side boundary of the CV, *x*^*^ =1, and correspond to the scenario with an infinite half-life of Aβ monomers and aggregates, 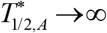 and 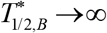, respectively. Results are shown for three values of the dimensionless diffusivity of Aβ monomers, 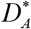.

The dimensionless concentrations of Aβ monomers and aggregates, 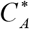 and 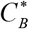, respectively, increase with the increase in 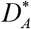, until 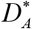 reaches a value of 10^6^ (corresponding to *D*_*A*_ of approximately 5 μm^2^/s for the values of *L* and *k*_1_ given in Table 3). It is worth noting that 5 μm^2^/s is an order of magnitude lower than the physiological value of 62.3 μm^2^/s given in Table 3. For values of 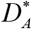 greater than this threshold, 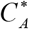 and 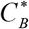 stabilize at a constant level, indicating that beyond this point, they are no longer constrained by diffusion (Fig. 2). In the scenario of dysfunctional protein degradation in neurons, with infinite half-lives for Aβ monomers and aggregates, the neurotoxic concentration of 50 μM (refer to Table 3) is reached when 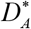 exceeds 10^5^, equivalent to *D*_*A*_ of around 0.5 µm^2^/s (which is two orders of magnitude less than the physiological value of 62.3 µm^2^/s). The approximate solutions for 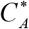 and 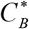 (given by Eqs. (S11) and (S10), respectively) do not depend on 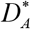. This is because the approximate solution is obtained under the lumped capacitance assumption, which assumes that 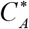 and 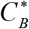 are independent of *x*^*^ and only depend on time *t*^*^. This corresponds to the assumption that 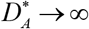. For this reason, the numerical solution computed for the scenario 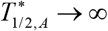 and 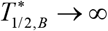 (the scenario for which the approximate solution was obtained) approaches the approximate solution for 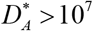 (Fig. 2).

A reduction in the dimensionless diffusivity, 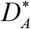, amplifies variations in 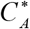 and 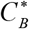 across the CV (Fig. S1). The numerical solution for a large value of 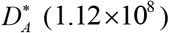 almost aligns with the approximate solution, represented by a horizontal line, as the approximate solution is unaffected by *x*^*^. The values of parameter ξ, as defined in Eqs. (14) and (20), are 0.263 (for 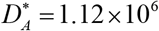), 0.0263 (for 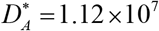), and 0.00263 (for 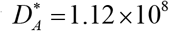). The decrease in parameter ξ (corresponding to larger values of 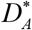) leads to the distributions of 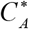 and 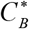 along *x*^*^ becoming closer to horizontal lines (Fig. S1). The independence of 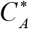 and 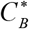 from *x*^*^ is necessary for the validity of the lumped capacitance approximation. It is noteworthy that the ratio of the variation of 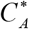 to the average value of 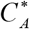 (Fig. S1a) is comparable to the ratio of the variation of 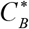 to the average value of 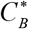 (Fig. S1b). This suggests that *ξ* ≪ 1 provides the criterion when both distributions of 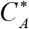 and 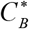 are close to uniform.

The decrease in 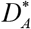 has minimal effect on 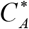 at the right-hand side boundary of the domain, *x*^*^ = 1 (Fig. S2a), and slightly decreases 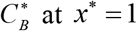, more noticeably for larger times. At 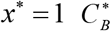 reaches the neurotoxic level at *t*^*^ = 0.5, approximately corresponding to a time, *t*, of 7 years (Fig. S2b). The numerical solutions for 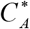 plotted in Fig. S2a closely resemble the approximate solution. In Fig. S2b, the numerical curves depicting 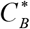 for larger values of 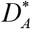 (1.12×10^7^ and 1.12×10^8^) coincide with the approximate solution. The unbounded linear growth of 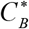 over time in Fig. S2b is attributed to the plot not accounting for the removal of Aβ aggregates from the cytosol due to deposition into Aβ plaques. The model assumes that all aggregates are eventually integrated into the plaques.

Over extended periods, the dimensionless radius of the Aβ plaque, 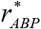, increases in proportion to the cube root of time, precisely in line with the approximate solution given by Eq. (27) (Fig. 3a). The independence of 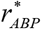 on Aβ monomer diffusivity is explained by the model assumption of a constant rate of Aβ monomer production, with all monomers entering the CV through the left-hand side boundary, and the assumption that all Aβ aggregates produced in the CV become part of the plaques. In the initial stages of plaque growth, when *t*^*^ is in the range [0 0.01], the growth of the plaque disagrees with the cube root of time behavior. The very beginning of the plaque growth is simulated by Eq. (29), although the plaque radius later deviates from Eq. (29) and depends on *t*^*^ in a sigmoidal manner until it matches the approximate solution (Fig. 3b). The fact that Eq. (29) describes the plaque growth only at very small times is not surprising since Eq. (29) only accounts for the effect of nucleation, described by kinetic constant *k*_1_. The sigmoidal shape of the numerically computed curve displaying 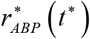 in the range of *t*^*^ between 0 and 0.01 is explained by autocatalytic growth, which takes over nucleation when enough nuclei are formed.

The independence of the plaque’s radius 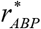 on 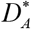 is confirmed in Fig. S3. The numerical solution coincides with the approximate solution for the scenario with 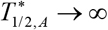 and 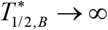.

### 3.3. Effects of varying the dimensionless rate of monomer production (the dimensionless monomer flux into the CV)

Effects of varying the dimensionless flux of Aβ monomers entering the CV through the left-hand side boundary, 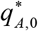, are investigated in Figs. 4, 5, and in Figs. S4-S6 in the Supplemental Materials.

**Fig. 4.**
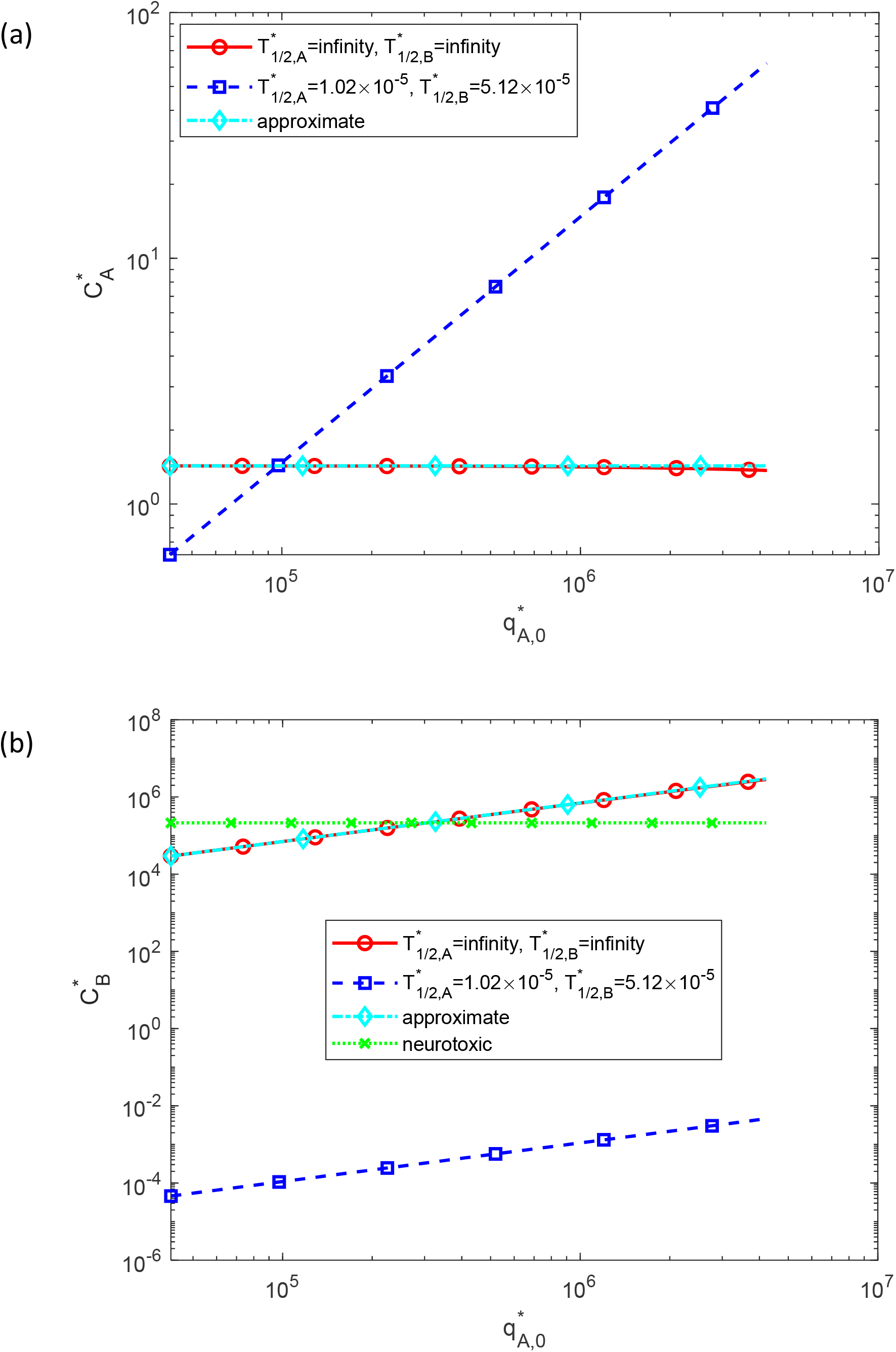
Dimensionless molar concentrations of Aβ monomers, 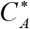 (a), and Aβ aggregates, 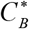 (b), plotted against the dimensionless flux of Aβ monomers from the left-hand side boundary of the CV, 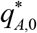. Approximate solutions for 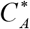 and 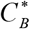 are computed using Eqs. (S11) and (S10), respectively. The approximate solution for 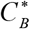 is directly proportional to 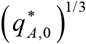. The computational results are presented at the right-hand side boundary of the CV, *x*^*^ =1, at 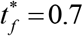 (10 years).

**Fig. 5.**
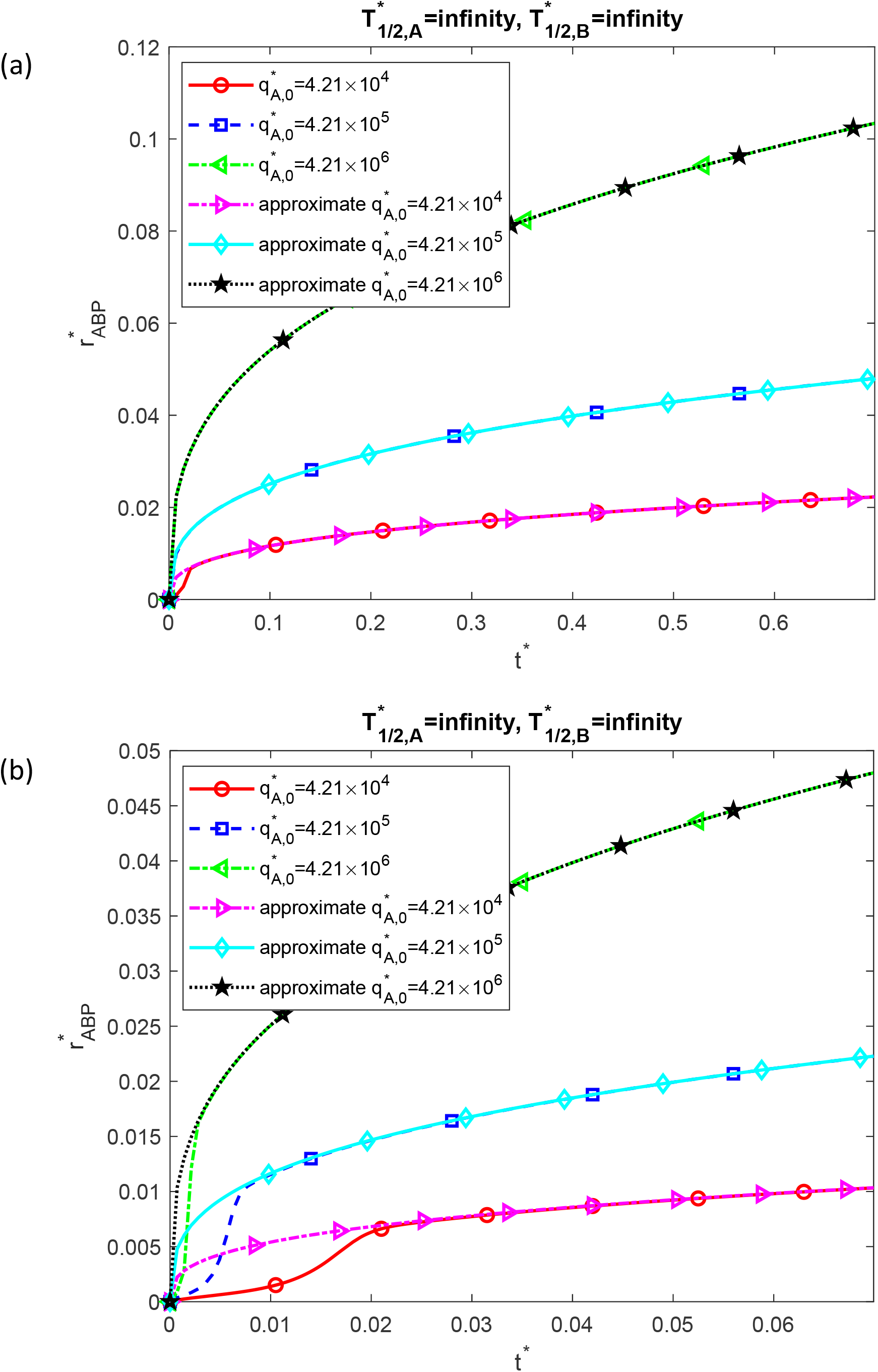
(a) Dimensionless radius of a growing Aβ plaque, 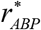, as a function of dimensionless time, *t*^*^. (b) Similar to Fig. 5a but focusing on the time range [0 0.07] on the *x*-axis. The approximate solution for 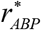 is plotted using Eq. (27); 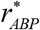 is directly proportional to 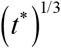. These computational results are presented at the right-hand side boundary of the CV, *x*^*^ =1, and correspond to the scenario with an infinite half-life of Aβ monomers and aggregates, 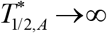 and 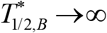, respectively. Results are shown for three values of the dimensionless flux of Aβ monomers from the left-hand side boundary of the CV, 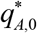.

The dimensionless concentration of Aβ monomers, 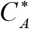, at *x*^*^ = 1 and 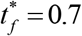 increases with 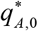 if the half-lives of Aβ monomers and aggregates are finite. If the half-lives are infinitely large, 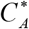 is independent of 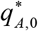 (extra monomers are converted into aggregates) (Fig. 4a). The approximate solutions for 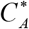 show little dependence on 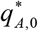. Indeed, the approximate solution for 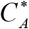 given by Eq. (S11) can be re-written as follows:

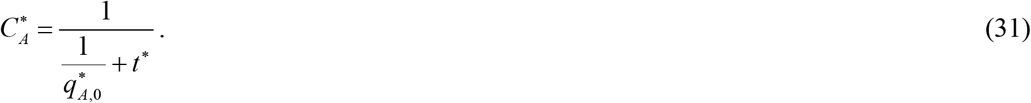

As 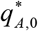 is large, the term 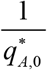 in the denominator of Eq. (31) is negligibly small. Fig. 4 is computed for *t*^*^ = 0.7. Hence, Eq. (31) gives 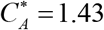, independent of 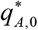 for large values of 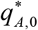. The numerical solution for 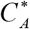 agrees with the approximate solution for the scenario with 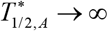 and 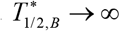 Fig. 4a).

The dimensionless concentration of Aβ aggregates, 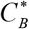, increases with the increase in 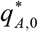 following a power relationship (depicted as a straight line in a log-log plot). For infinite half-lives of monomers and aggregates, the neurotoxic concentration of aggregates is reached when 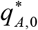 exceeds 3×10^5^, equivalent to *q*_*A*,0_ of approximately 8×10^−6^ μM μm s^−1^ (Fig. 4b). The approximate solution for 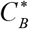 is directly proportional to 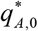 (refer to Eq. (S10)). The curves depicting the numerical and approximate solutions coincide for the scenario with 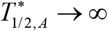 and 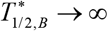.

An increase in the dimensionless flux of Aβ monomers, 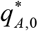, increases the variation in 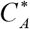 across the CV (Fig. S4a). As discussed above, the approximate solution for 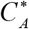 exhibits a weak dependence on 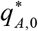. The numerical solution for 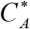 agrees with the approximate solution for a small value of 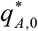 (4.21×10^4^). However, for larger values of 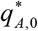 (4.21×10^5^ and 4.21×10^6^), a deviation from the lumped capacitance conditions occurs, and 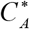 displays variations across the CV, not captured by the approximate solution. The increase in 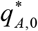 also shifts the average value of 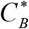 in the CV to greater values and increases the variation in 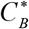 across the CV (Fig. S4b). The numerical and approximate solutions for 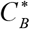 coincide for smaller values of 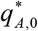 (4.21×10^4^ and 4.21×10^5^), while for 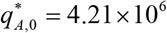 the numerical solution displays a variation across the CV, deviating from the approximate solution. The parameter *ξ*, as defined in Eqs. (14) and (20), takes values of 0.00263 (for 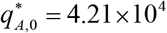), 0.0263 (for 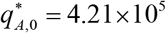), and 0.263 (for 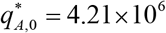). A decrease in *ξ* (corresponding to smaller values of 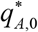) causes the distributions of 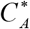 and 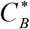 along *x*^*^ to approach horizontal lines (Fig. S4). Therefore, reducing ξ aligns the distributions of 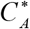 and 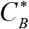 along *x*^*^ closer to those predicted by the lumped capacitance approximation.

The dimensionless concentration of Aβ monomers, 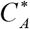, at the right-hand side boundary of the domain, *x*^*^ = 1, exhibits an initial rapid increase followed by a gradual decrease over time (Fig. S5a). The numerical and approximate solutions in Fig. S5a nearly coincide. The dimensionless concentration of Aβ aggregates, 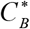, at *x*^*^ = 1 exhibits a linear relationship with time (Fig. S5b). The neurotoxic threshold is reached faster with higher values of the monomer flux. For example, for 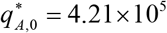 the aggregate concentration 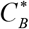 reaches the neurotoxic level at *t*^*^ = 0.52, which approximately corresponds to *t* of 7.5 years while for 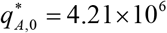 the aggregate concentration, 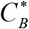, reaches the neurotoxic level at *t*^*^ = 0.06, which approximately corresponds to *t* of 1 year (Fig. S5b). The numerical and approximate solutions agree for smaller values of 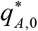 (4.21×10^4^ and 4.21×10^5^), but some deviation is observed for 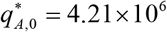 due to the breakdown of lumped capacitance conditions.

The dimensionless radius of the Aβ plaque, 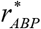, increases proportionally to the cube root of time, in precise agreement with the approximate solution given by Eq. (27) (Fig. 5a). Note that according to Eq. (27), 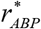 is proportional to 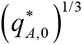. Hence, the results for different values of 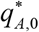 in Fig. 5a are different. However, for each value of 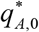, the numerical and analytical solutions coincide at large times. In contrast, for small times (when *t*^*^ falls within the range [0 0.02] for 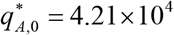, [0 0.01] for 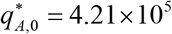, and [0 0.005] for 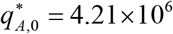), the numerical solution deviates from the analytical one. In these small time ranges the numerical solution displays a sigmoidal increase in the plaque’s radius, consistent with observations in Burgold et al. (2014).

Confirmation of the power relationship between 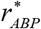 and 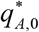 is presented in Fig. S6 (with the power on 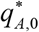 being 1/3). The numerical and approximate solutions in Fig. S6 coincide for the scenario with 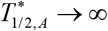 and 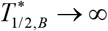.

### 3.4. Effects of varying the dimensionless half-life of Aβ monomers

Effects of varying the dimensionless half-life of Aβ monomers, 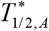, are investigated in Figs. 6, 7, and in Figs. S7-S9 in the Supplemental Materials.

**Fig. 6.**
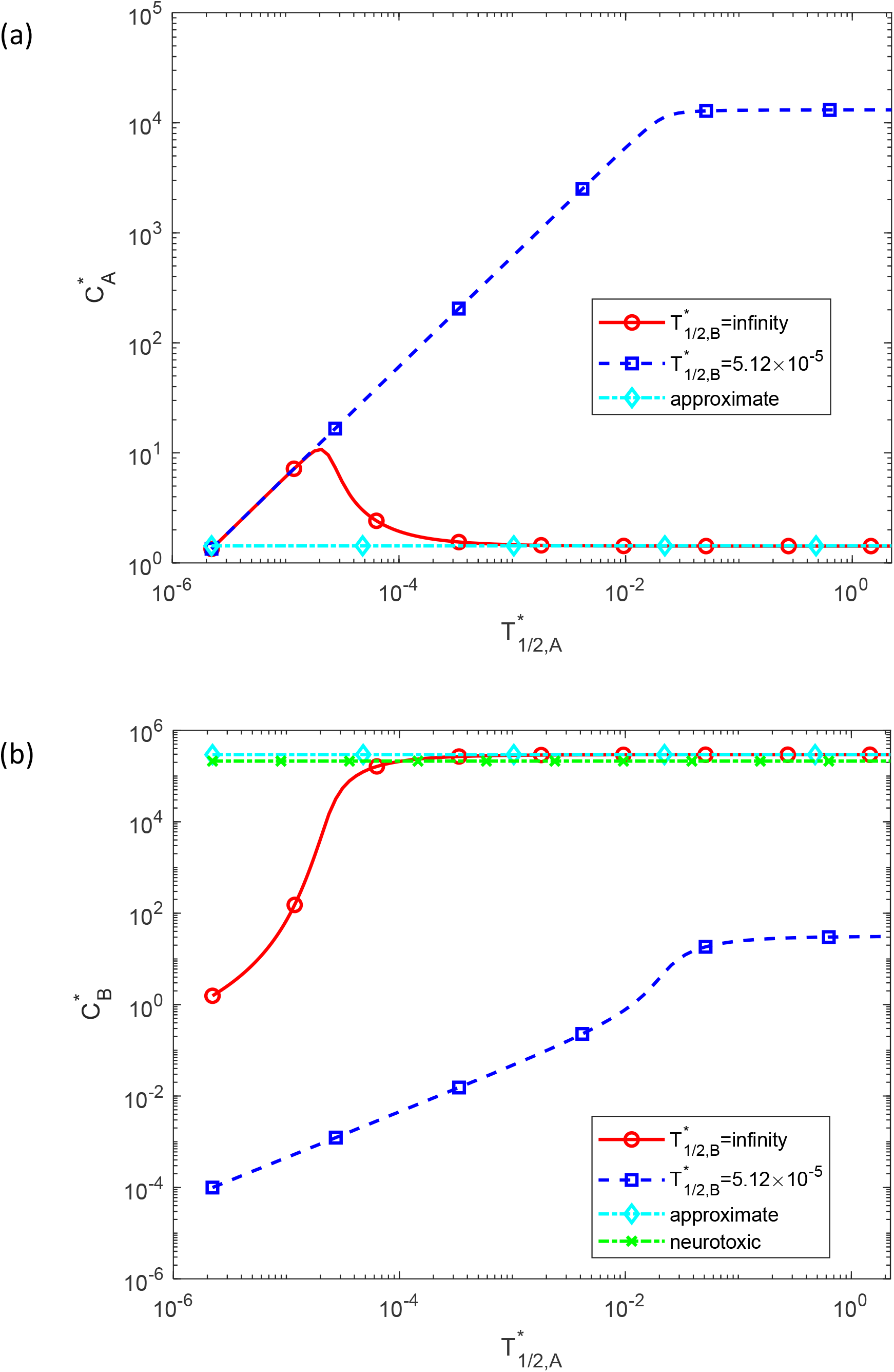
Dimensionless molar concentration of Aβ monomers, 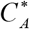 (a), and Aβ aggregates, 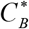 (b), plotted against the dimensionless half-life of Aβ monomers, 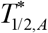. Approximate solutions for 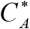 and 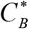 are computed using Eqs. (S11) and (S10), respectively. The computational results are presented at the right-hand side boundary of the CV, *x*^*^ =1, at 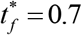 (10 years).

**Fig. 7.**
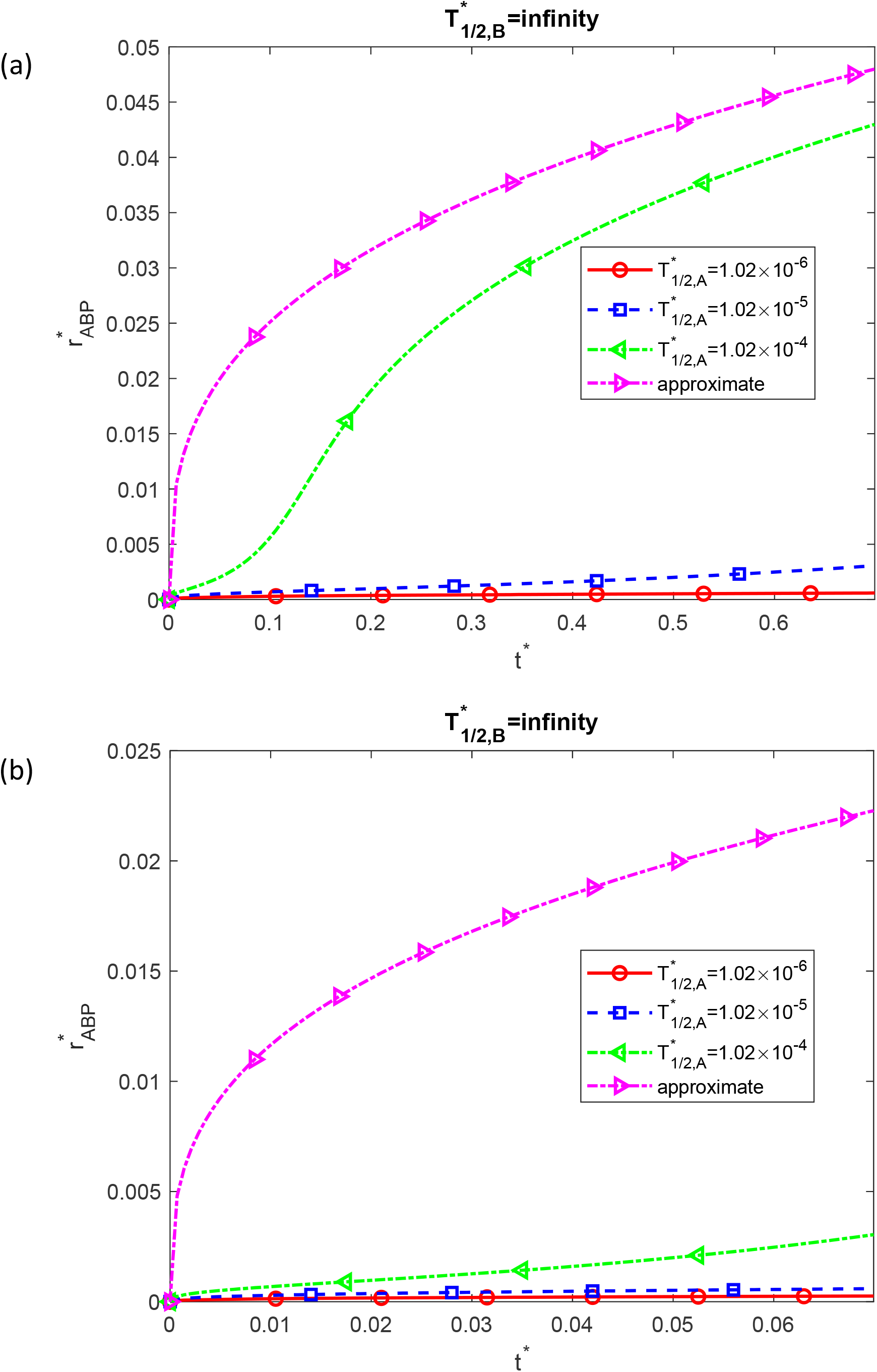
(a) Dimensionless radius of a growing Aβ plaque, 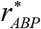, as a function of dimensionless time, *t*^*^. (b) Similar to Fig. 7a but focusing on the time range [0 0.07] on the *x*-axis. The approximate solution for 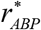 is plotted using Eq. (27); 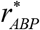 is directly proportional to (*t*^*^)^1/3^. These computational results are presented at the right-hand side boundary of the CV, *x*^*^ =1, and correspond to the scenario with an infinite half-life of Aβ aggregates, 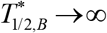. Results are shown for three values of the dimensionless half-life of Aβ monomers, 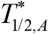.

The dimensionless concentration of Aβ monomers, 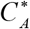, at *x*^*^ = 1 and 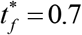 increases as 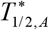 increases until 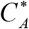 reaches a constant value, for the case with a finite half-life of Aβ aggregates, 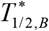. In contrast, for the case with the infinite half-life of Aβ aggregates, 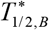, 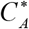 initially increases as 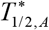 increases and then decreases to a constant value (Fig. 6a). The dimensionless concentration of Aβ aggregates, 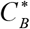, increases as 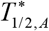 increases until reaching a constant value. In the infinite half-life of Aβ aggregates scenario, the neurotoxic concentration of aggregates is reached when 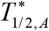 exceeds 10^−4^, equivalent to a *T*_1/ 2, *A*_ of around 5×10^4^ s (Fig. 6b). The approximate solution is independent of the values of 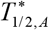 and 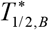 (because it is obtained under the assumption that 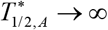 and 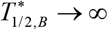). Numerical solutions for 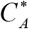 and 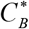 obtained for 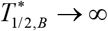 approach the respective approximate solutions when 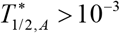 in Fig. 6a and 6b.

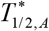 has no effect on variations of 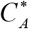 and 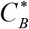 across the CV (Fig. S7). Since the approximate solution is derived for 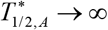 and 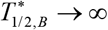, there is a slight discrepancy between the numerical and approximate solutions for 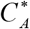 and 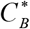, even for the largest value of 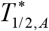 (1.02 ×10^−4^) (Fig. S7).

The effect of 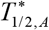 on the dimensionless concentration of Aβ monomers, 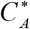, at the right-hand side boundary of the domain, *x*^*^ = 1, depends on the value of 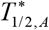. For large 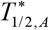 (1.02×10^−4^), 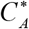 exhibits a rapid increase from zero (a value that is dictated by the initial condition in Eq. (17a)) followed by a gradual decrease over time. Conversely, for smaller 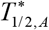 (1.02×10^−5^ and 1.02×10^−6^), 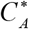 quickly increases from zero to a constant value, with the constant value being smaller for a smaller value of 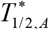 (Fig. S8a). The effect of 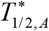 on the dimensionless concentration of Aβ aggregates, 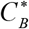, at the right-hand side boundary of the domain, *x*^*^ = 1, also depends on the value of 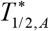. A large value of 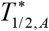 (1.02×10^−4^) results in an increase in 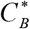 until 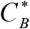 reaches a neurotoxic concentration at *t*^*^ = 0.7, corresponding to 10 years. However, for smaller values of 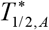 (1.02×10^−5^ and 1.02×10^−6^), 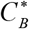 remains close to zero (Fig. S8b). Some discrepancy between the numerical and approximate solutions in Fig. S8, even for the largest value of 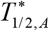 (1.02×10^−4^), is attributed to the fact that this value of 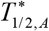 is still insufficiently close to infinity.

The dimensionless radius of the Aβ plaque, 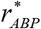, increases as dimensionless time, *t* ^*^, increases, yet even for the largest value of 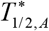 (1.02×10^−4^) it remains below the curve depicting the approximate solution (27) (Fig. 7a). The validity of the approximate solution is limited to infinitely long half-lives of Aβ monomers and aggregates. As the decay of monomers results in less conversion of monomers into aggregates, it leads to fewer aggregates being incorporated into the plaques. At the rates of Aβ monomer degradation used to compute Fig. 7, the plaque grows slowly at small *t*^*^, slower for smaller values of 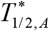 (Fig. 7b).

Fig. S9 demonstrates that for the case of an infinite half-life of Aβ aggregates, the dimensionless plaque radius reaches that predicted by the approximate solution when 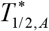 exceeds 10^−3^, which approximately corresponds to 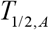 of 4.5×10^5^ s.

### 3.5. Effects of varying the dimensionless half-life of Aβ aggregates

Effects of varying the dimensionless half-life of Aβ aggregates, 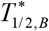, are investigated in Figs. 8, 9, and in Figs. S10-S13 in the Supplemental Materials.

**Fig. 8.**
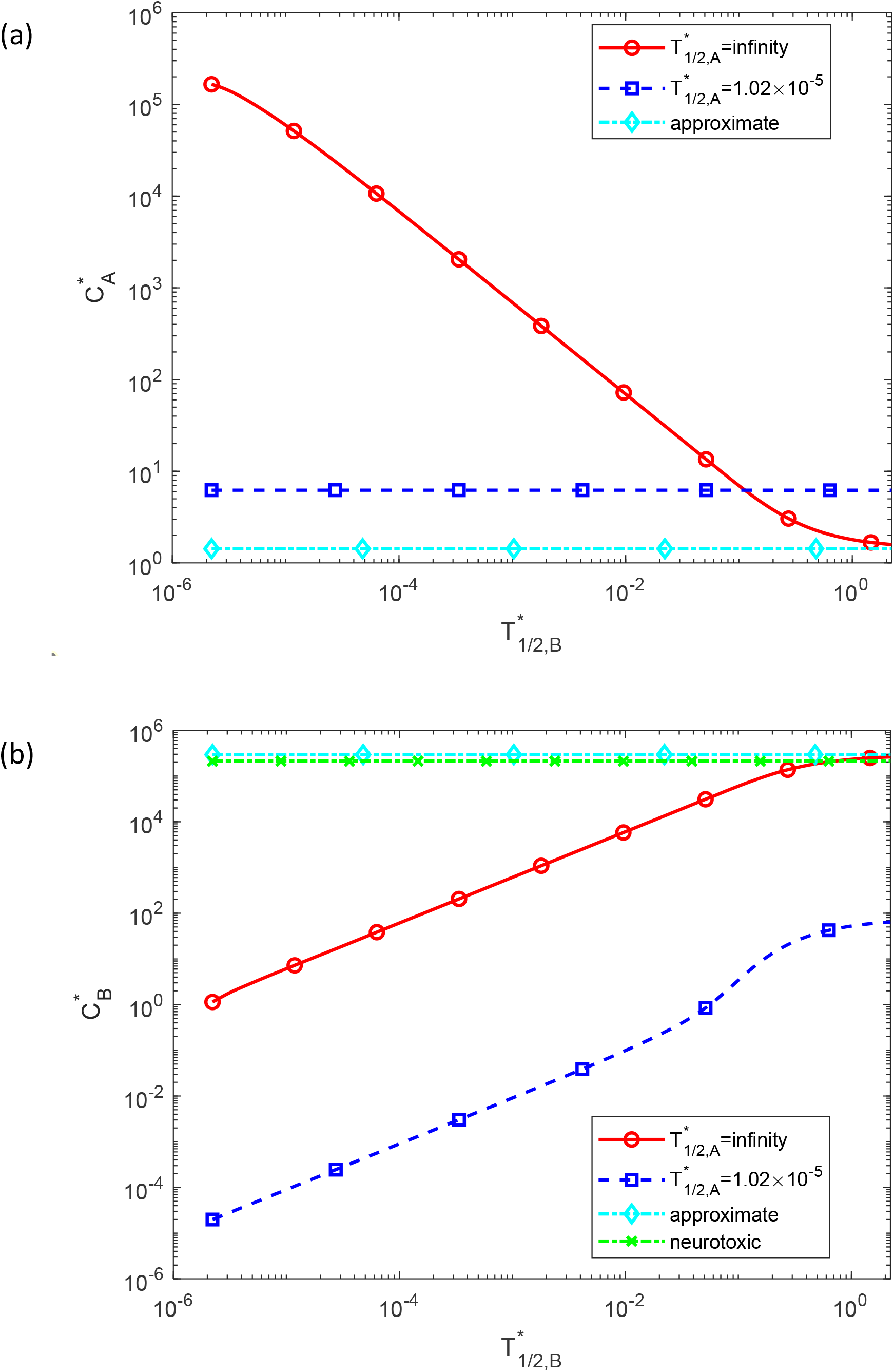
Dimensionless molar concentration of Aβ monomers, 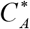 (a), and Aβ aggregates, 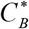 (b), plotted against the dimensionless half-life of Aβ aggregates, 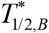. Approximate solutions for 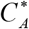 and 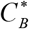 are computed using Eqs. (S11) and (S10), respectively. The computational results are presented at the right-hand side boundary of the CV, *x*^*^ =1, at 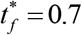 (10 years).

**Fig. 9.**
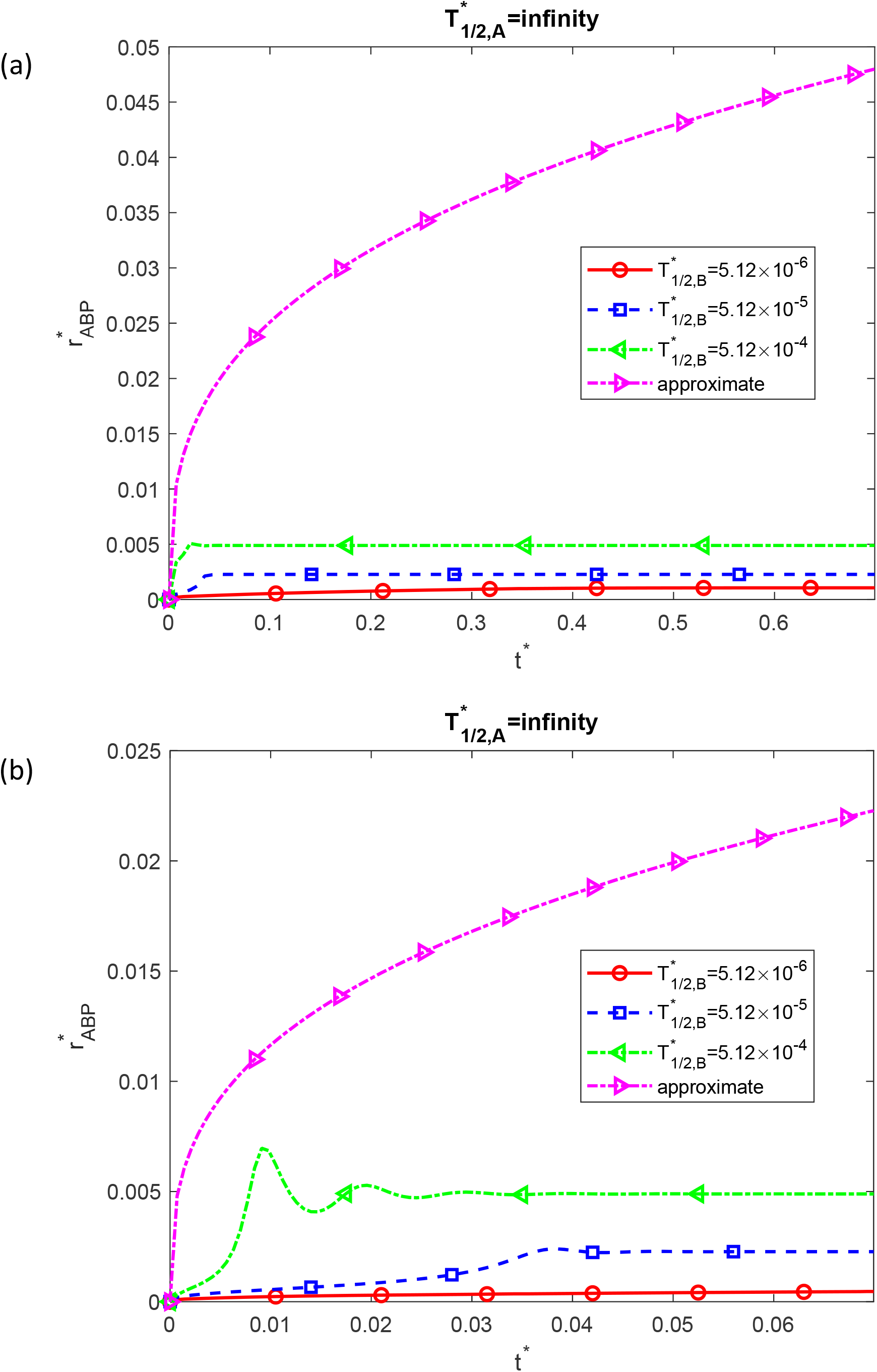
(a) Dimensionless radius of a growing Aβ plaque, 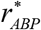, as a function of dimensionless time, *t*^*^. (b) Similar to Fig. 9a but focusing on the time range [0 0.07] on the *x*-axis. The approximate solution for 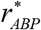 is plotted using Eq. (27); 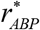 is directly proportional to (*t*^*^)^1/3^. These computational results are presented at the right-hand side boundary of the CV, *x*^*^ =1, and correspond to the scenario with an infinite half-life of Aβ monomers, 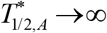. Results are shown for three values of the dimensionless half-life of Aβ aggregates, 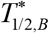.

The dimensionless concentration of Aβ monomers, 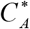, at *x*^*^ = 1 and 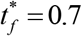 decreases with the increase in 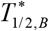, for the case with the infinite half-life of Aβ monomers, 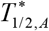. However, for the scenario with a finite half-life of Aβ monomers, 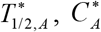 remains constant (Fig. 8a). The dimensionless concentration of Aβ aggregates, 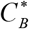, increases as 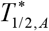 increases. In the case of an infinite half-life of Aβ monomers, the neurotoxic concentration of aggregates is reached when 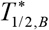 exceeds 1, approximately corresponding to *T*_1/ 2, *B*_ of around 4.5×10^8^ s (Fig. 8b). In Fig. 8, both 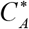 and 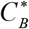 approach the approximate solution for the scenario with 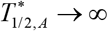 when 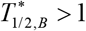.

The parameter 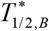 has no effect on variations in 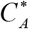 and 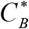 across the CV (Fig. S10). The dimensionless concentration of Aβ monomers, 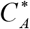, at the right-hand side boundary of the domain, *x*^*^ = 1, gradually increases with time until it reaches a constant value, which is of Aβ aggregates, 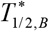 (Fig. S11a). The dimensionless concentration of Aβ aggregates, 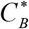, at the right-hand side boundary of the domain, *x*^*^ = 1, remains much smaller than the neurotoxic concentration. It also stays significantly below the curve representing the approximate solution for 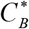, even for the largest value of 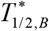 (5.12×10^−4^) (Fig. S11b).

The dimensionless radius of the Aβ plaque, 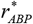, increases with an increase in the dimensionless time, *t* ^*^. However, even for the largest value of 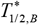 (5.12×10^−4^), it remains much below the curve predicted by the approximate solution given by Eq. (27) (Fig. 9a). Interestingly, for small times, the curve depicting the case with 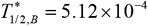 oscillates in the range of *t* ^*^ between 0 and 0.03 (Fig. 9b). To verify that this is not a numerical artifact, computations were repeated on a mesh ten times finer with respect to *t* ^*^ and *x*^*^. The result displayed in Fig. S12 shows an oscillatory behavior identical to that displayed in Fig. 9b.

Additionally, Fig. S13 shows that for the case of an infinite half-life of Aβ monomers, the dimensionless plaque radius reaches the value predicted by the approximate solution when 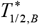 exceeds 2.2, corresponding to *T*_1/ 2,*B*_ of 10^9^ s.

## 4. Discussion, model limitations, and future directions

This study introduces a model that simulates the diffusion of Aβ monomers, the conversion of monomers into aggregates, and the formation of senile plaques. The density of Aβ plaque is assumed to be 1.35 g/cm^3^, and the growth time of an Aβ plaque is set at 10 years. Under these conditions the model involves four dimensionless parameters: (1) the dimensionless diffusivity of Aβ monomers, (2) the dimensionless flux of Aβ monomers into the CV due to monomer production on lipid membranes, (3) the dimensionless half-life of Aβ monomers, and (4) the dimensionless half-life of Aβ aggregates. It is demonstrated that the validity of the lumped capacitance approximation for this problem depends on the smallness of a new dimensionless parameter. This parameter is defined as the product of the dimensionless flux of Aβ monomers into the CV and the dimensionless duration of the process, divided by the dimensionless diffusivity of Aβ monomers.

Given the nonlinear nature of the governing equations, the relationships between the solutions and the parameters are complex, as anticipated for nonlinear problems. Key observations and insights are summarized below.

The guidance on how the solution depends on the dimensionless parameters is provided by the approximate solution. The approximate solution is given by Eq. (S10), providing the concentration of aggregates, 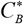; Eq. (S11), providing the concentration of monomers, 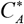; and Eq. (27), specifying the plaque radius, 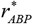. The approximate solution indicates that, over extended periods, aggregate concentration linearly increases with *t*^*^, while monomer concentration decreases inversely proportional to *t*^*^. 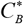 is directly proportional to the monomer flux, 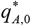, while 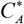 is independent of 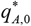 for large *t*^*^. These trends are observed in Figs. 4 and S5. The plaque radius is proportional to the cube root of time. For a fixed density *ρ*^*^, 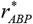 only depends on the flux of monomers into the CV, 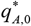; it is proportional to 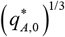. These trends are observed in Figs. 5 and S6.

However, the approximate solution has limitations stemming from two assumptions used to obtain it: (i) the lumped capacitance approximation, where variations in concentrations of Aβ monomers and aggregates across the CV are considered negligible, and concentrations depend solely on time, and (ii) the assumption of dysfunctional protein degradation machinery, implying infinitely large half-lives for Aβ monomers and aggregates.

The lumped capacitance model fails when the diffusivity of Aβ monomers is low (Figs. 2 and S1). Notably, for the lumped capacitance model to break down due to small diffusivity of Aβ monomers, the diffusivity must be an order of magnitude less than its physiological value. In essence, with the physiological value of the monomer diffusivity, Aβ plaque formation is not diffusion-controlled. The lumped capacitance model may also fail for very large fluxes of Aβ monomers into the CV (Fig. S4).

The assumption of infinite half-lives for Aβ monomers and aggregates is more restrictive. Reducing 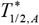 and 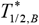 from infinite to physiological values results in significant deviations from the approximate solution (Figs. 6-9). For finite half-lives of Aβ monomers and/or aggregates, the Aβ plaque radius does not grow indefinitely with time, which agrees with the existence of the saturation state reported in Serrano-Pozo et al. (2012). The plaque radius stabilizes once it reaches a certain value dependent on the half-life of monomers/aggregates (Fig. 9). This is consistent with Serrano-Pozo et al. (2012) whose results suggest that during the pathological progression of Alzheimer’s disease, plaques attain their maximum size early on and experience minimal growth thereafter. For finite half-lives of monomers and aggregates, the solution’s dependence on parameters becomes more complex, involving all four parameters: 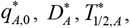, and 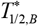 (while the approximate solution depends only on 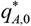, see Eqs. (S10), (S11), and (27)).

The model developed in this study has limitations as it does not simulate the process of Aβ aggregate assembly into plaques. Instead, the model assumes that all aggregates produced in the CV are absorbed by the plaque. Thus, calculating the rate of production of aggregates makes it possible to simulate the growth of the plaque. Future research should focus on simulating the assembly of aggregates into plaques. This would allow for improving the accuracy of the model because Aβ aggregates exhibit autocatalytic properties (see Eqs. (3) and (4)), and thus the removal of aggregates into the plaques is expected to reduce the rate of monomer conversion into aggregates. It is important to investigate scenarios where new plaques deposit close to existing ones, forming plaque clusters as suggested in McCarter et al. (2013). Additionally, simulating the age-related increase in the number of plaques (McCarter et al., 2013) is a crucial avenue for future exploration.

## Abbreviations

Aβ: amyloid beta
AD: Alzheimer’s disease
APP: amyloid precursor protein
CV: control volume
F-W: Finke-Watzky
PDE: partial differential equation

## Data accessibility

This article has no additional data.

## Authors’ contributions

AVK is the sole author of this paper.

## Supplemental Materials

### S1. Exact and approximate solutions of Eqs. (15)-(19) for the case of dysfunctional protein degradation machinery

The approximate and exact solutions for Eqs. (15)-(19) were derived in Kuznetsov (2024). The procedure is summarized below.

If Aβ aggregation were diffusion-limited, aggregate growth would predominantly occur at the tips, resulting in a branching, tree-like structure. The absence of such structures implies that the process is not diffusion-limited (Cruz et al., 1997).

For a large diffusivity of Aβ monomers, the CV can be treated as a lumped capacitance body, where concentrations depend solely on time, not on location. In this case, Eqs. (15)-(19) can be reformulated as the following system of ordinary differential equations:

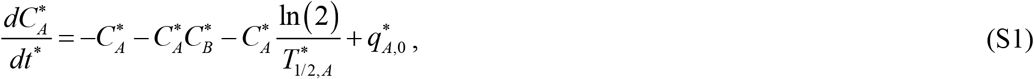

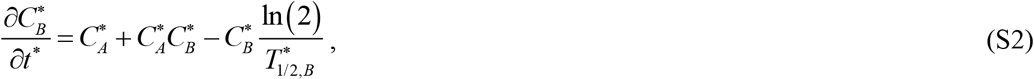

which must be solved subject to the following initial conditions:

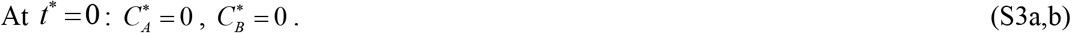

The exact and approximate solutions are derived for the scenario when the protein degradation machinery is dysfunctional, implying infinitely large half-lives of Aβ monomers and aggregates, 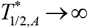 and 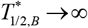. When Eqs. (S1) and (S2) are added for this scenario, the following result is obtained:

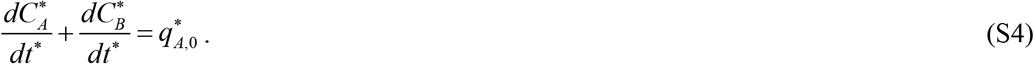

By integrating Eq. (S4) and using the initial conditions given by Eq. (S3), the following result is obtained:

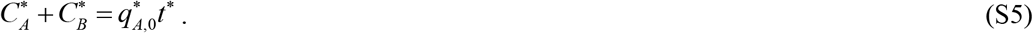

The increase in 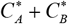 over time is attributed to the continuous production of Aβ monomers. These monomers undergo continuous conversion into aggregates, and in the assumed scenario characterized by 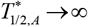 and 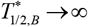, neither monomers nor aggregates are subject to destruction.

The exact solution was derived using the following approach. 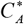 is eliminated from Eq. (S2) through the utilization of Eq. (S5):

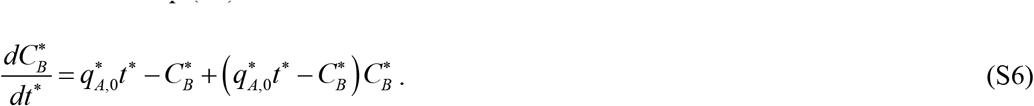

Eq. (S6) is similar to the one examined in Kuznetsov and Kuznetsov (2022). To find the exact solution of Eq. (S6) with the initial condition given by Eq. (S3b), the DSolve function, followed by the FullSimplify function in Mathematica 13.3 (Wolfram Research, Champaign, IL), was utilized. The resulting solution is presented below:

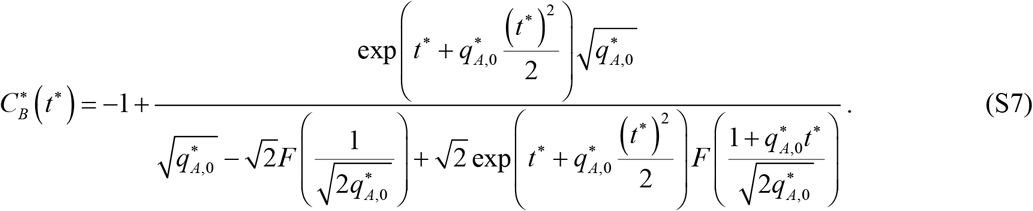

In this equation, *F* (*x*) is Dawson’s integral:

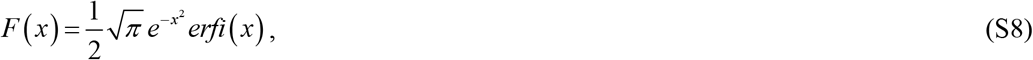

where *erfi* (*x*) represents the imaginary error function.

If *t*^*^ → 0, Eq. (S7) implies that

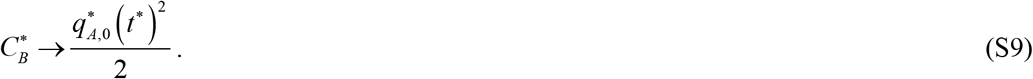

As the exact solution (S7) is hard to analyze for large *t*^*^ (due to the NaN error), an approximate solution valid for large values of *t*^*^ is derived. The analysis of numerical solutions in Kuznetsov (2024) indicates that as time increases, the concentration of Aβ aggregates linearly increases while the concentration of Aβ monomers decreases. Substituting this result into Eq. (S5) yields:

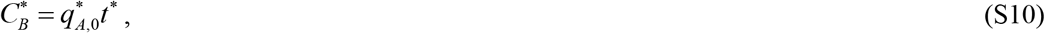

which is valid for large *t*^*^.

Substituting this into Eq. (S2), the following is obtained:

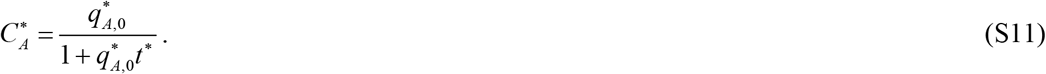

Eqs. (S10) and (S11) constitute the approximate solution for the concentrations of Aβ aggregates and monomers, respectively. Note that 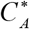 given by Eq. (S11) violates the initial condition given by Eq. (17a) and hence is valid only for large *t*^*^.

In dimensional variables, Eq. (S11) can be rewritten as:

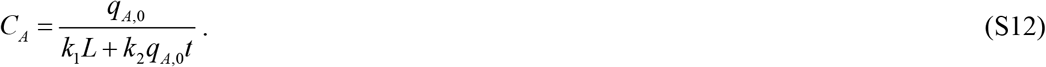

### S2. Supplementary figures

**Fig. S1.**
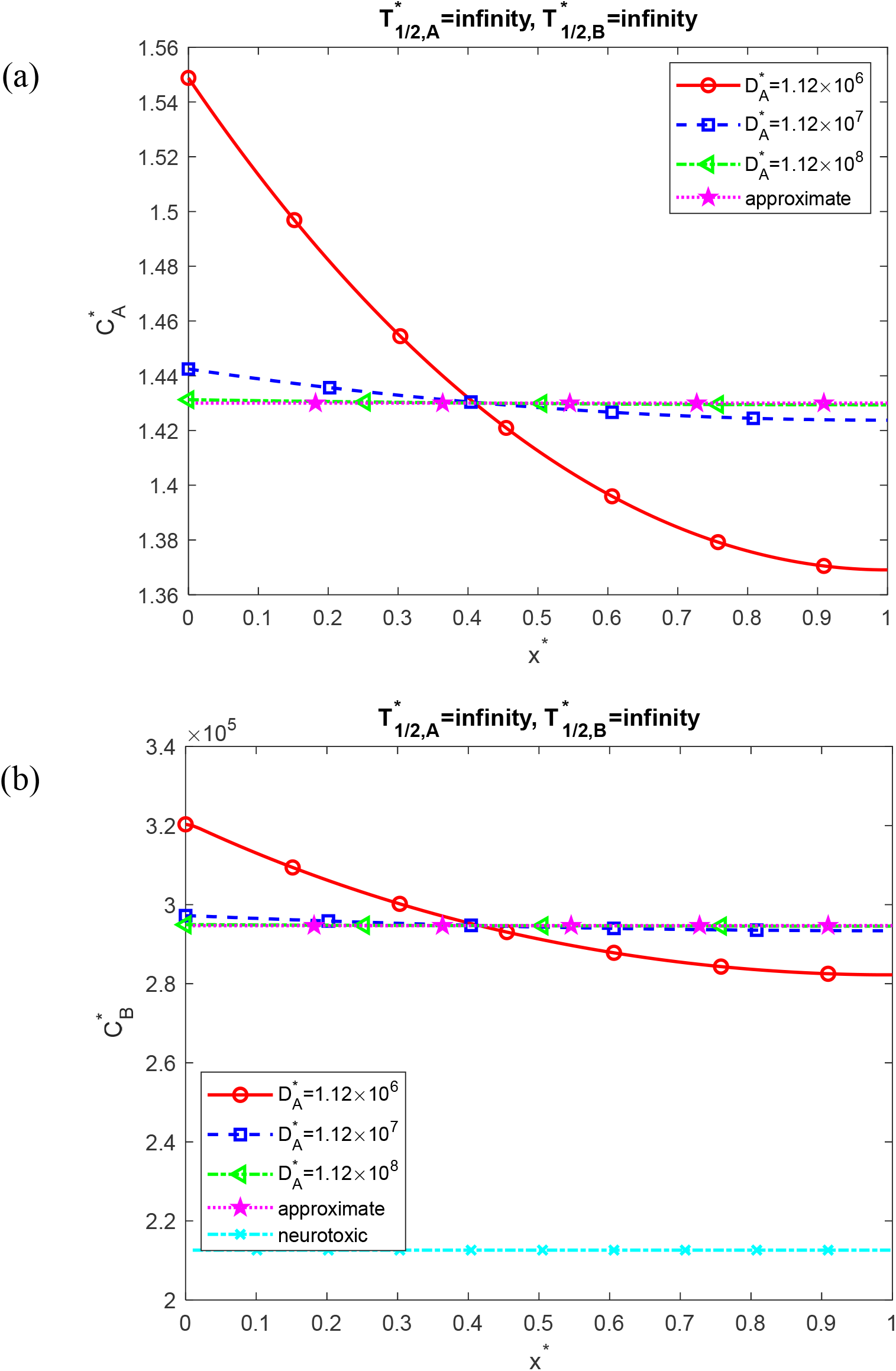
Dimensionless molar concentrations of Aβ monomers, 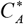 (a) and Aβ aggregates, 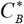 (b) as a function of the dimensionless distance from the surface releasing Aβ monomers (e.g., cell membrane), *x*^*^. Approximate solutions (independent of *x*^*^) for 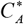 and 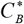 are computed using Eqs. (S11) and (S10), respectively. These results are presented at 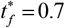 (10 years) and correspond to the scenario with an infinite half-life of Aβ monomers and aggregates, 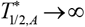 and 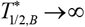, respectively. Results are shown for three values of the dimensionless diffusivity of Aβ monomers, 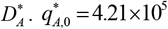.

**Fig. S2.**
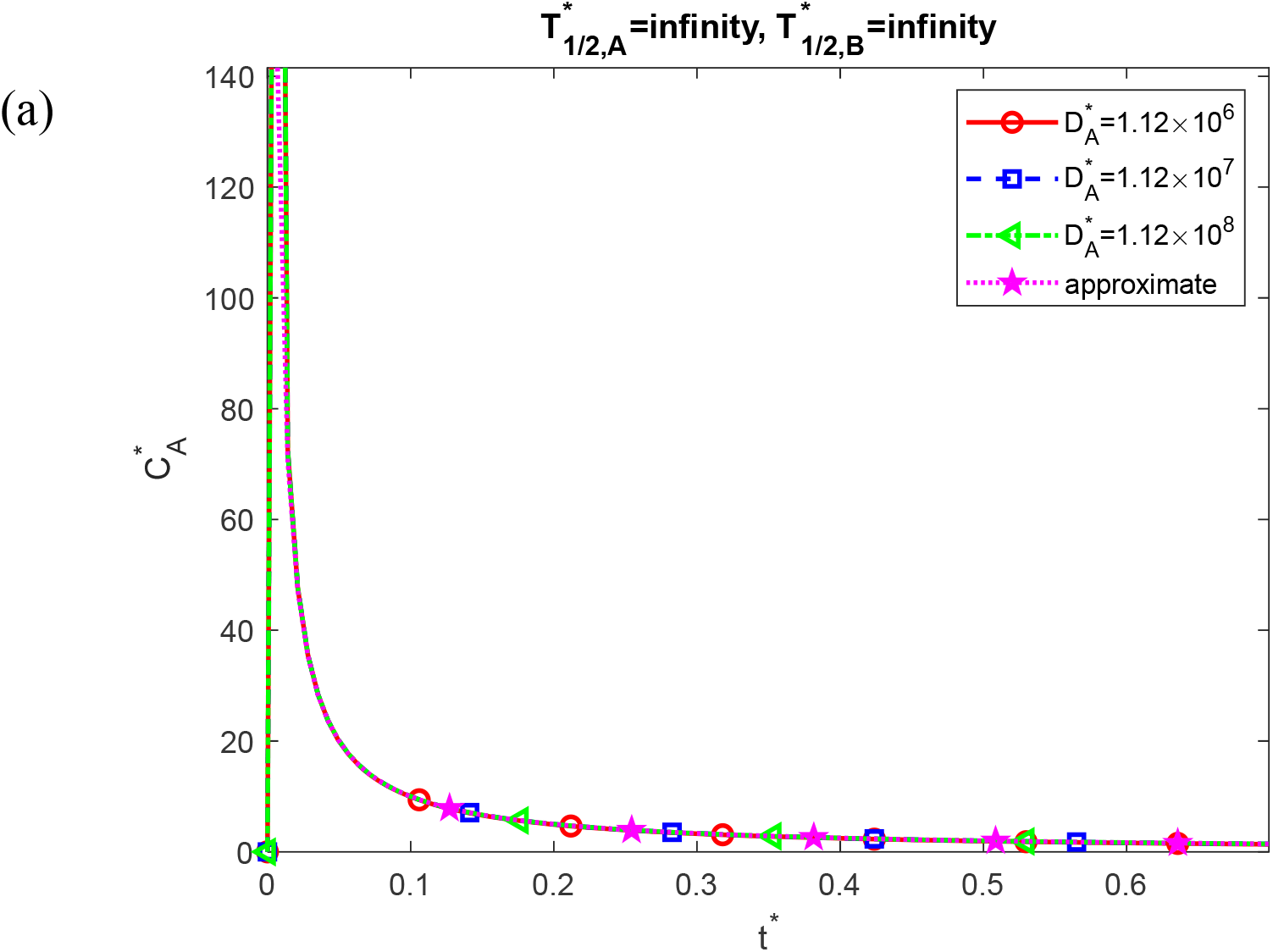
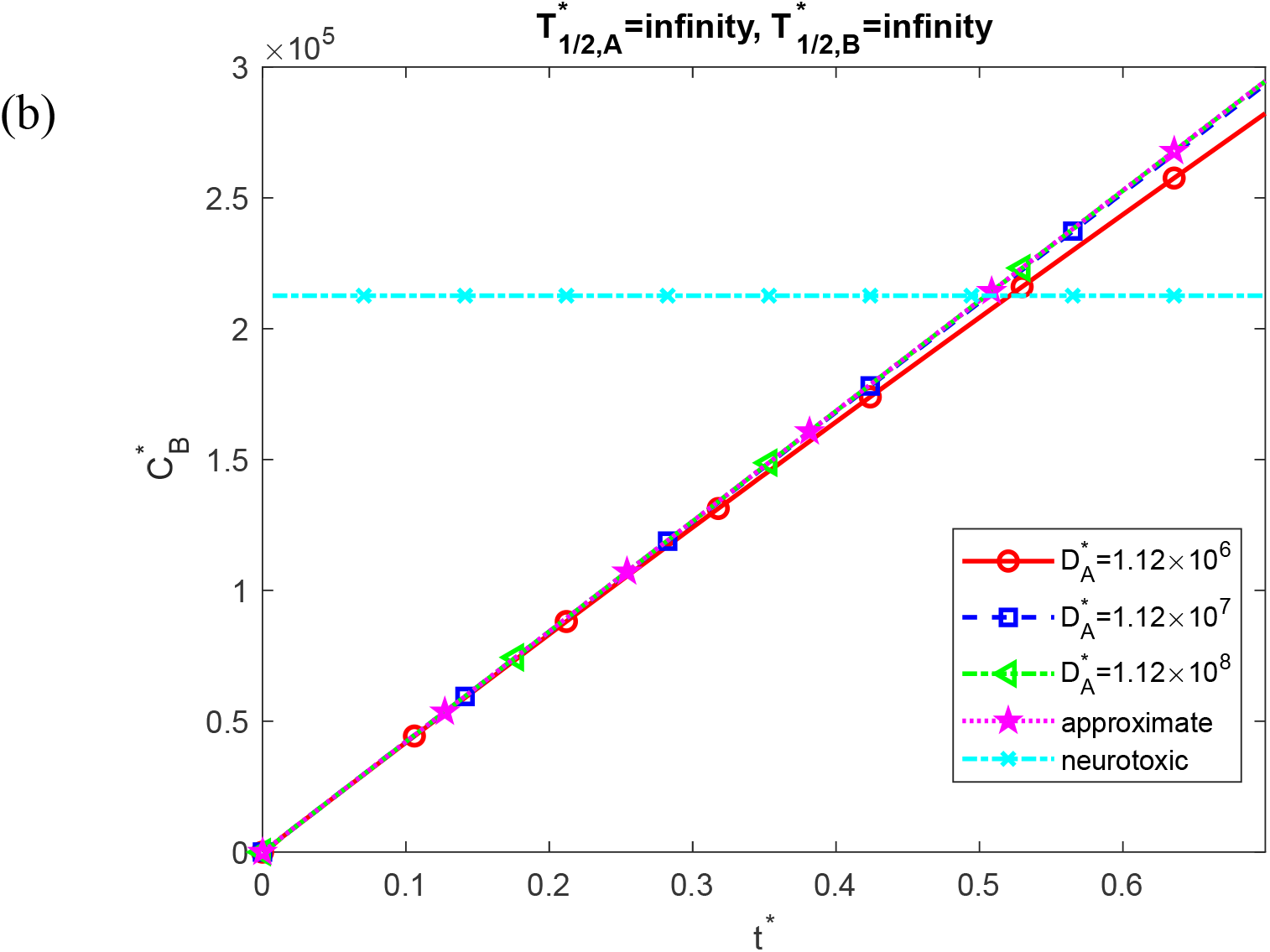
Dimensionless molar concentrations of Aβ monomers, 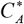 (a) and Aβ aggregates, 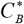 (b) as a function of dimensionless time, *t*^*^. Approximate solutions for 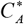 and 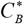 are computed using Eqs. (S11) and (S10), respectively. The approximate solution for 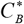 is directly proportional to *t*^*^. These results are presented at the right-hand side boundary of the CV, *x*^*^ =1, and correspond to the scenario with an infinite half-life of Aβ monomers and aggregates, 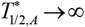 and 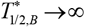, respectively. Results are shown for three values of the dimensionless diffusivity of Aβ monomers, 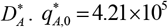.

**Fig. S3.**
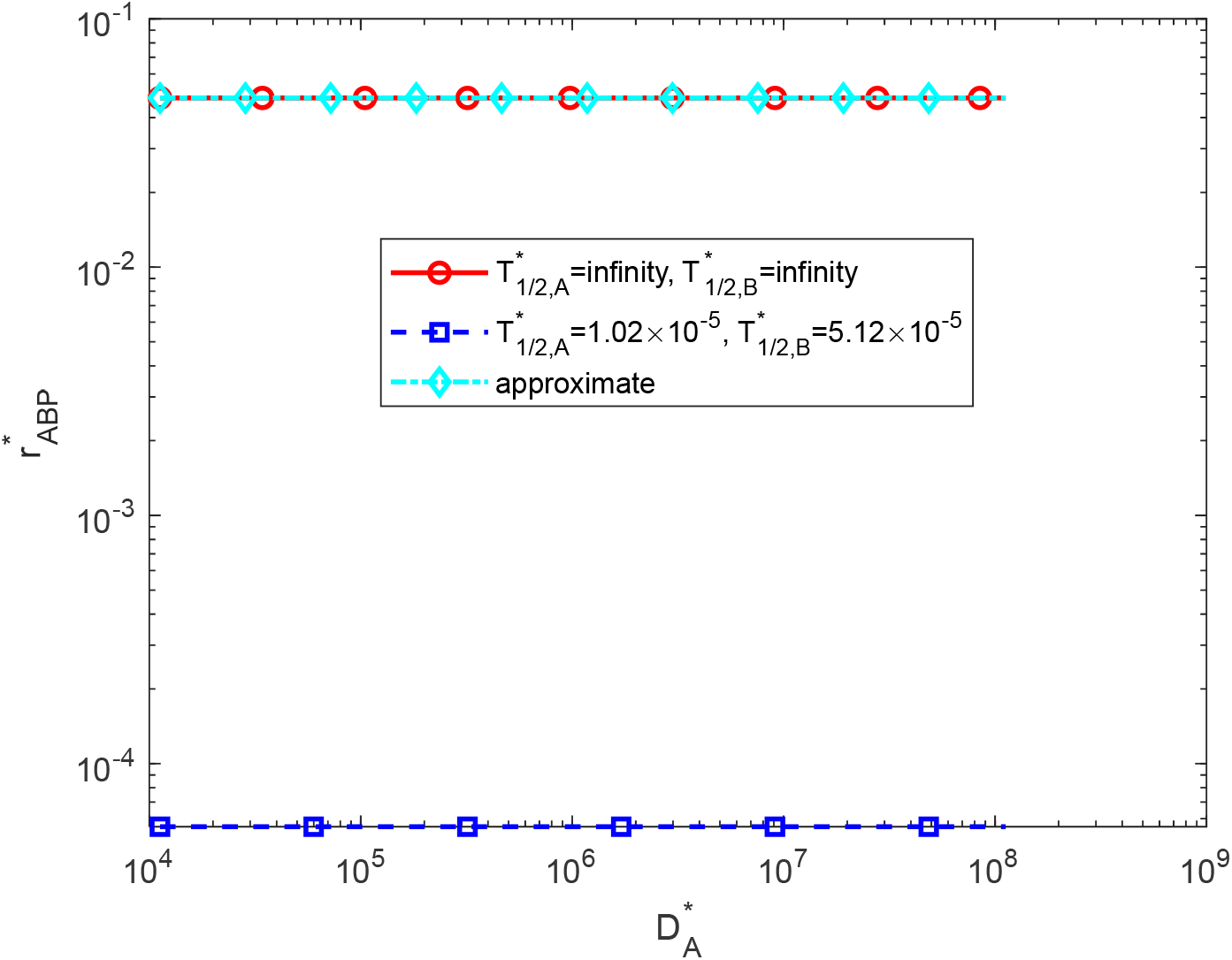
Dimensionless radius of an Aβ growing plaque, 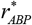, as a function of the dimensionless diffusivity of Aβ monomers, 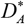, at 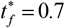 (10 years). The dimensionless radius of Aβ plaque for infinite half-life of Aβ monomers and aggregates is 0.0480 (corresponding dimensional radius is 2.40 μm). 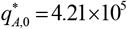.

**Fig. S4.**
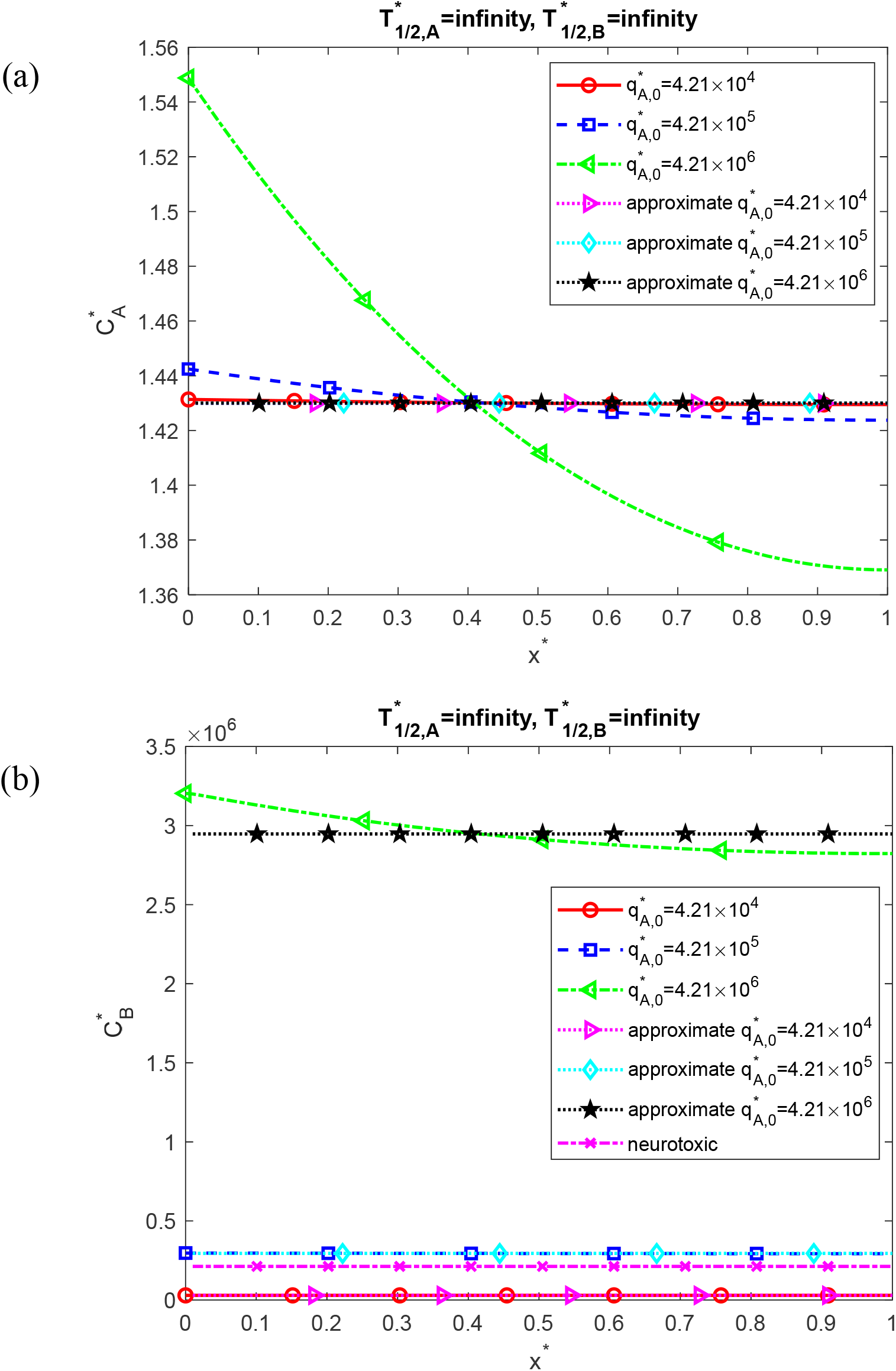
Dimensionless molar concentrations of Aβ monomers, 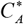 (a) and Aβ aggregates, 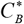 (b) as a function of the dimensionless distance from the surface releasing Aβ monomers (e.g., cell membrane), *x*^*^. Approximate solutions (independent of *x*^*^) for 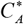 and 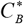 are computed using Eqs. (S11) and (S10), respectively. These results are presented at 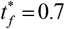 (10 years) and correspond to the scenario with an infinite half-life of Aβ monomers and aggregates, 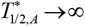 and 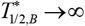, respectively. Results are shown for three values of the dimensionless flux of Aβ monomers from the left-hand side boundary of the CV, 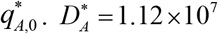.

**Fig. S5.**
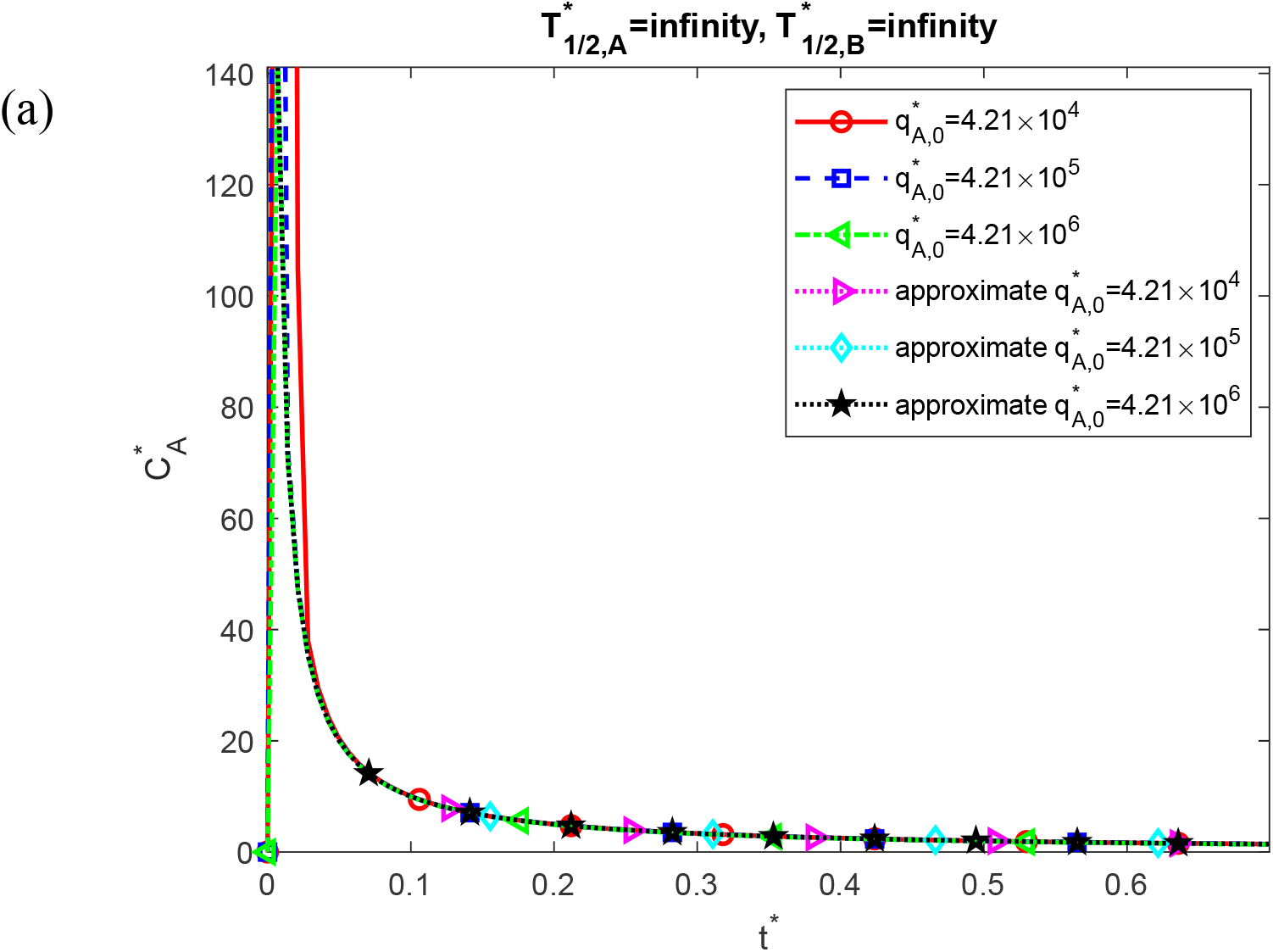
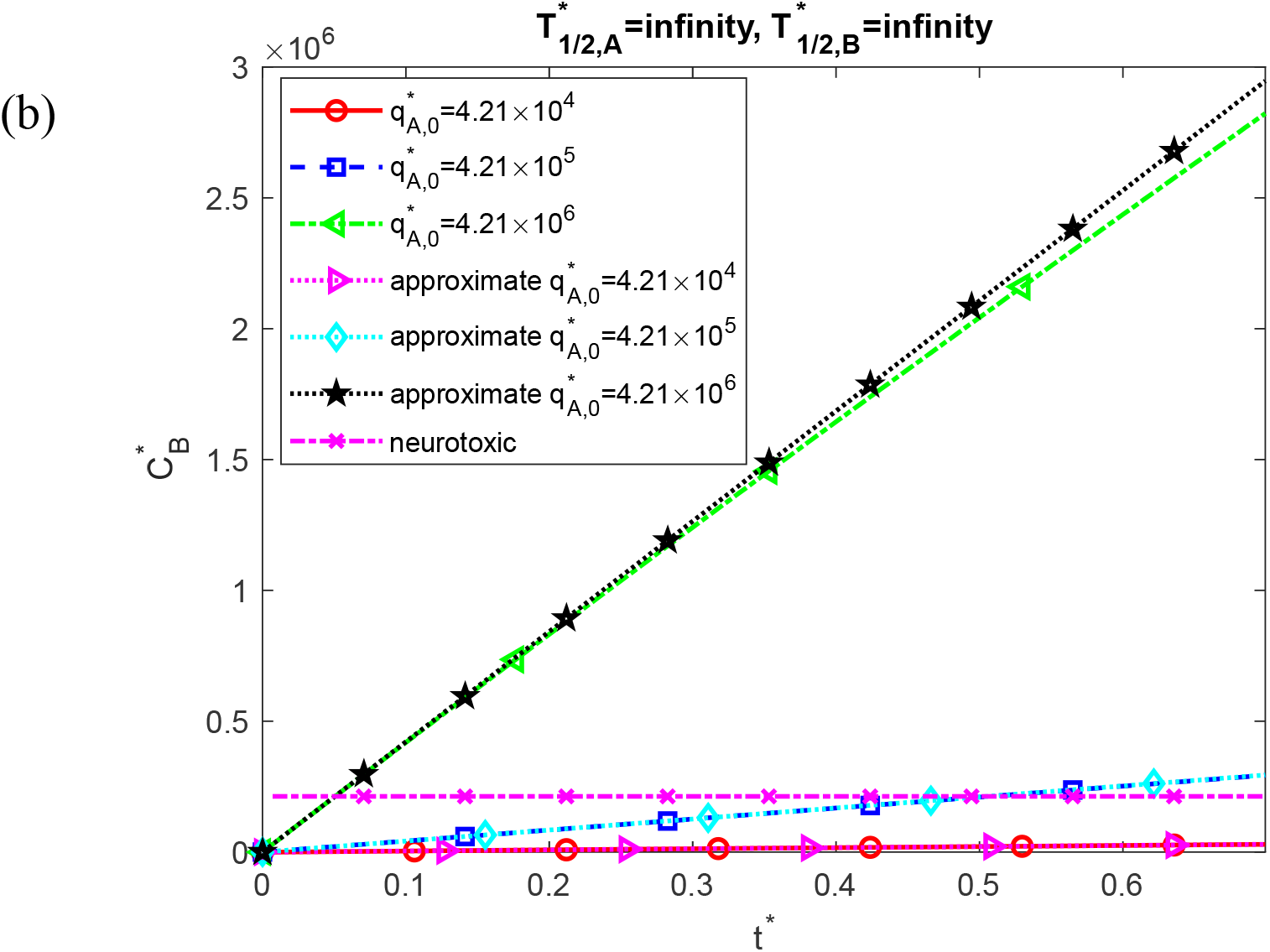
Dimensionless molar concentrations of Aβ monomers, 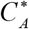 (a) and Aβ aggregates, 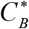 (b) as a function of dimensionless time, *t*^*^. Approximate solutions for 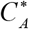 and 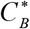 are computed using Eqs. (S11) and (S10), respectively. The approximate solution for 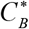 is directly proportional to *t*^*^. These results are presented at the right-hand side boundary of the CV, *x*^*^ =1, and correspond to the scenario with an infinite half-life of Aβ monomers and aggregates, 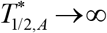 and 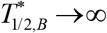, respectively. Results are shown for three values of the dimensionless flux of Aβ monomers from the left-hand side boundary of the CV, 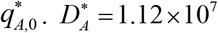.

**Fig. S6.**
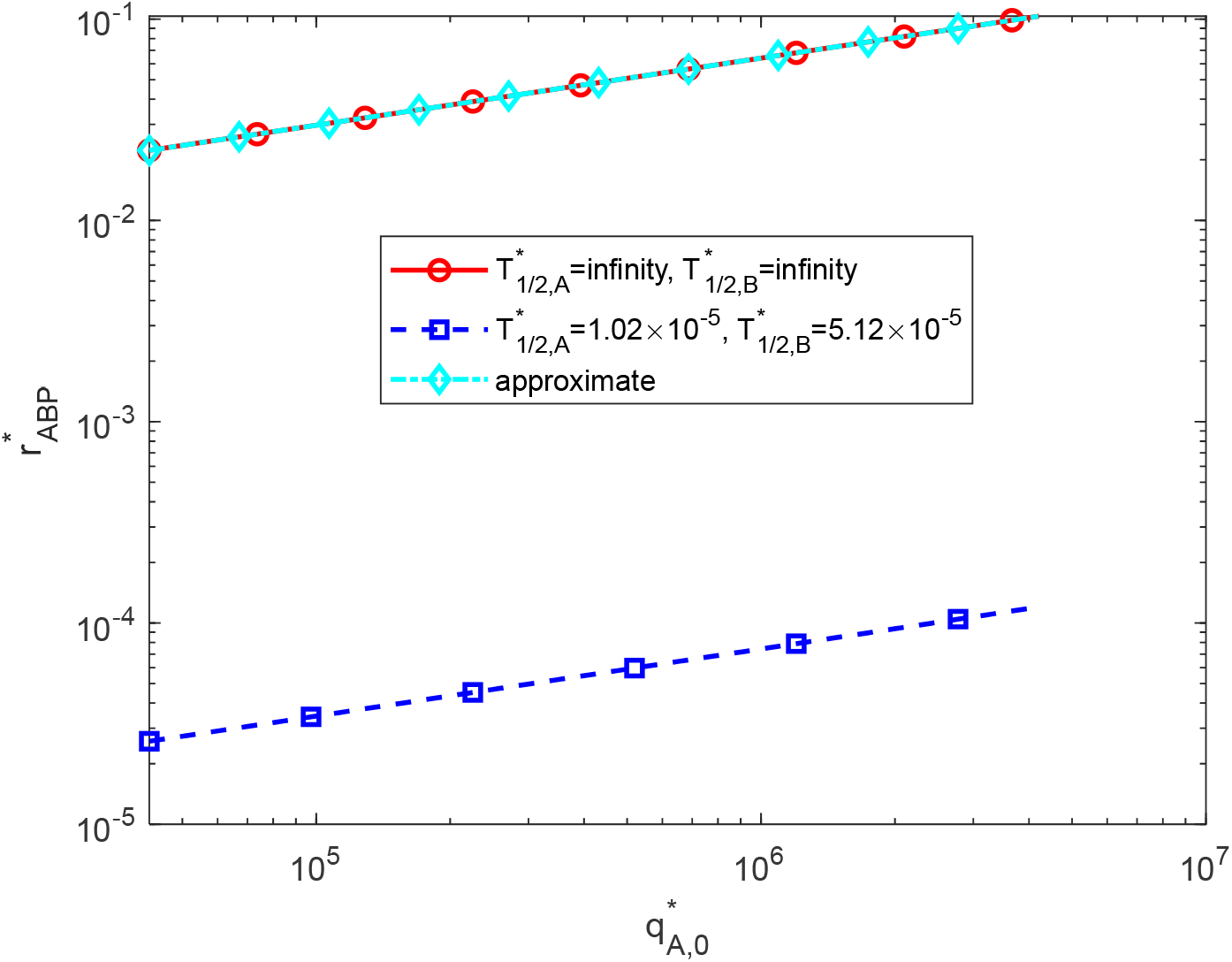
Dimensionless radius of a growing Aβ plaque, 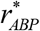, as a function of the dimensionless flux of Aβ monomers from the left-hand side boundary of the CV, 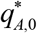, at 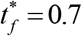 (10 years). Dimensionless radius of Aβ plaque corresponding to the largest value of *q*^*^ in Fig. S6 is 0.103 (corresponding dimensional radius is 5.17 μm); the result shown is for the infinite half-life of Aβ monomers and aggregates. 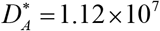.

**Fig. S7.**
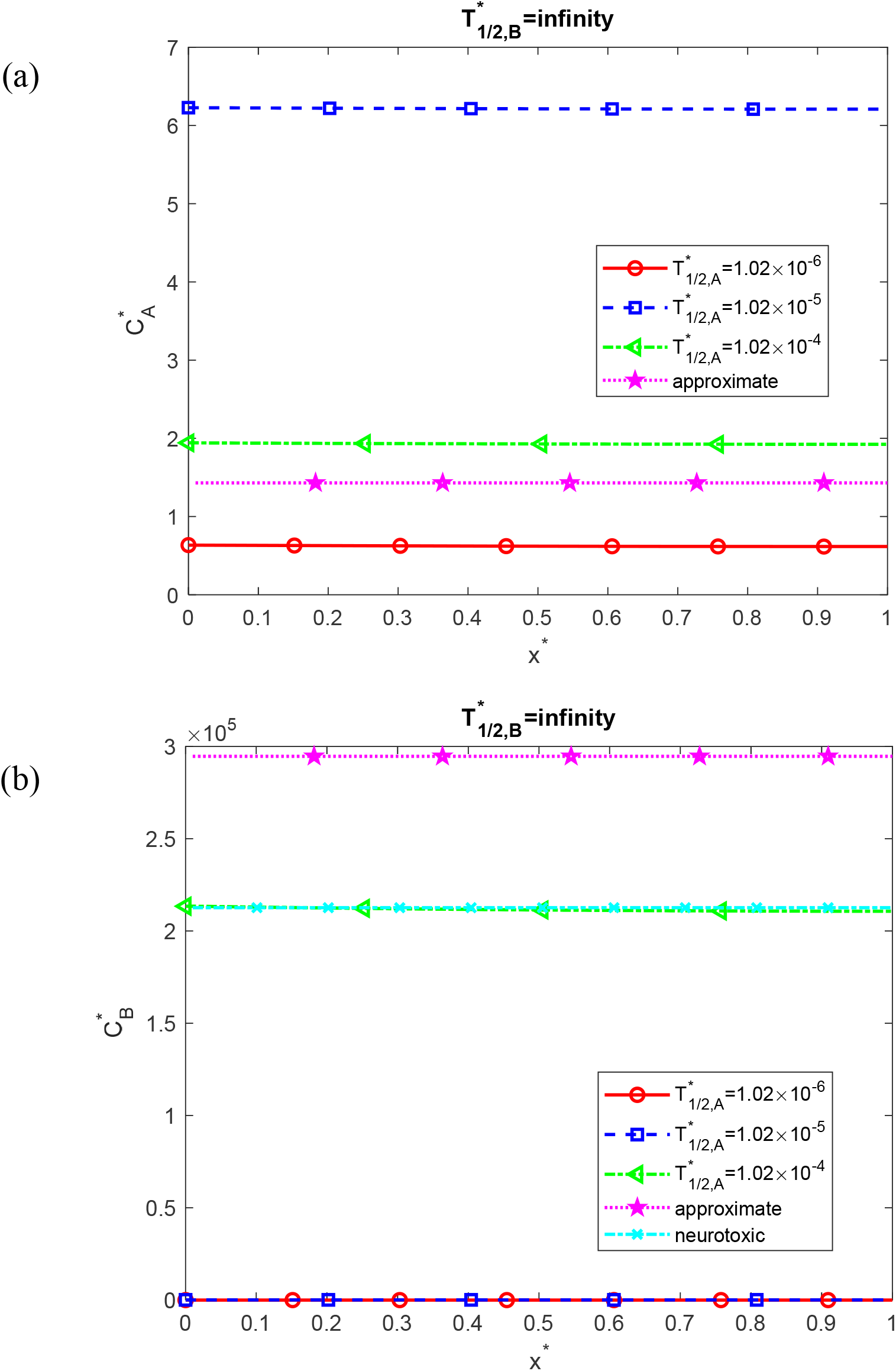
Dimensionless molar concentrations of Aβ monomers, 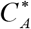 (a) and Aβ aggregates, 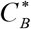 (b) as a function of the dimensionless distance from the surface releasing Aβ monomers (e.g., cell membrane), *x*^*^. Approximate solutions (independent of *x*^*^) for 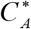 and 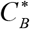 are computed using Eqs. (S11) and (S10), respectively. These results are presented at 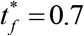 (10 years) and correspond to the scenario with an infinite half-life of Aβ aggregates, 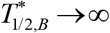. Results are shown for three values of the dimensionless half-life of Aβ monomers, 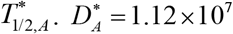 and 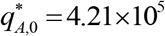.

**Fig. S8.**
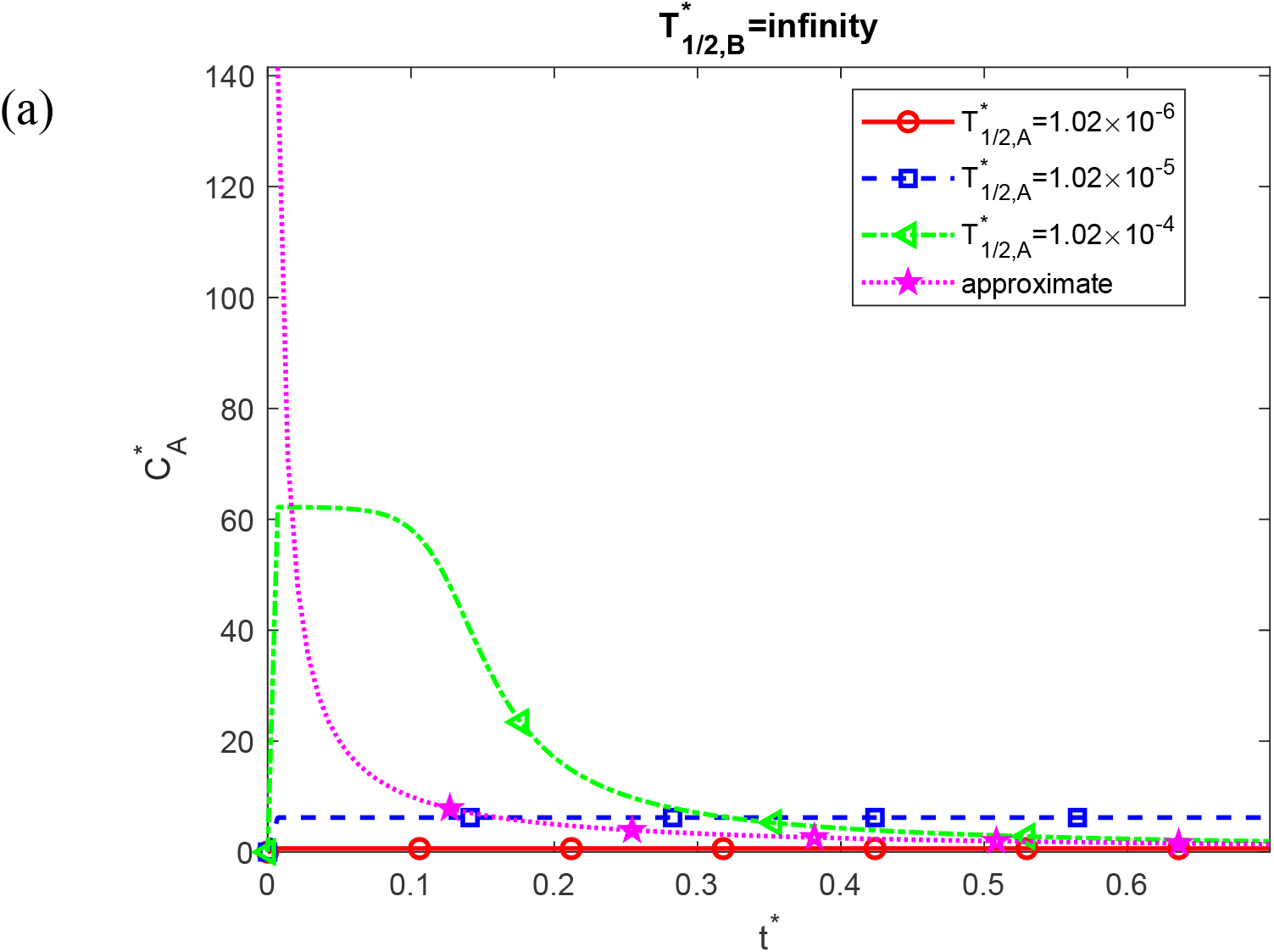
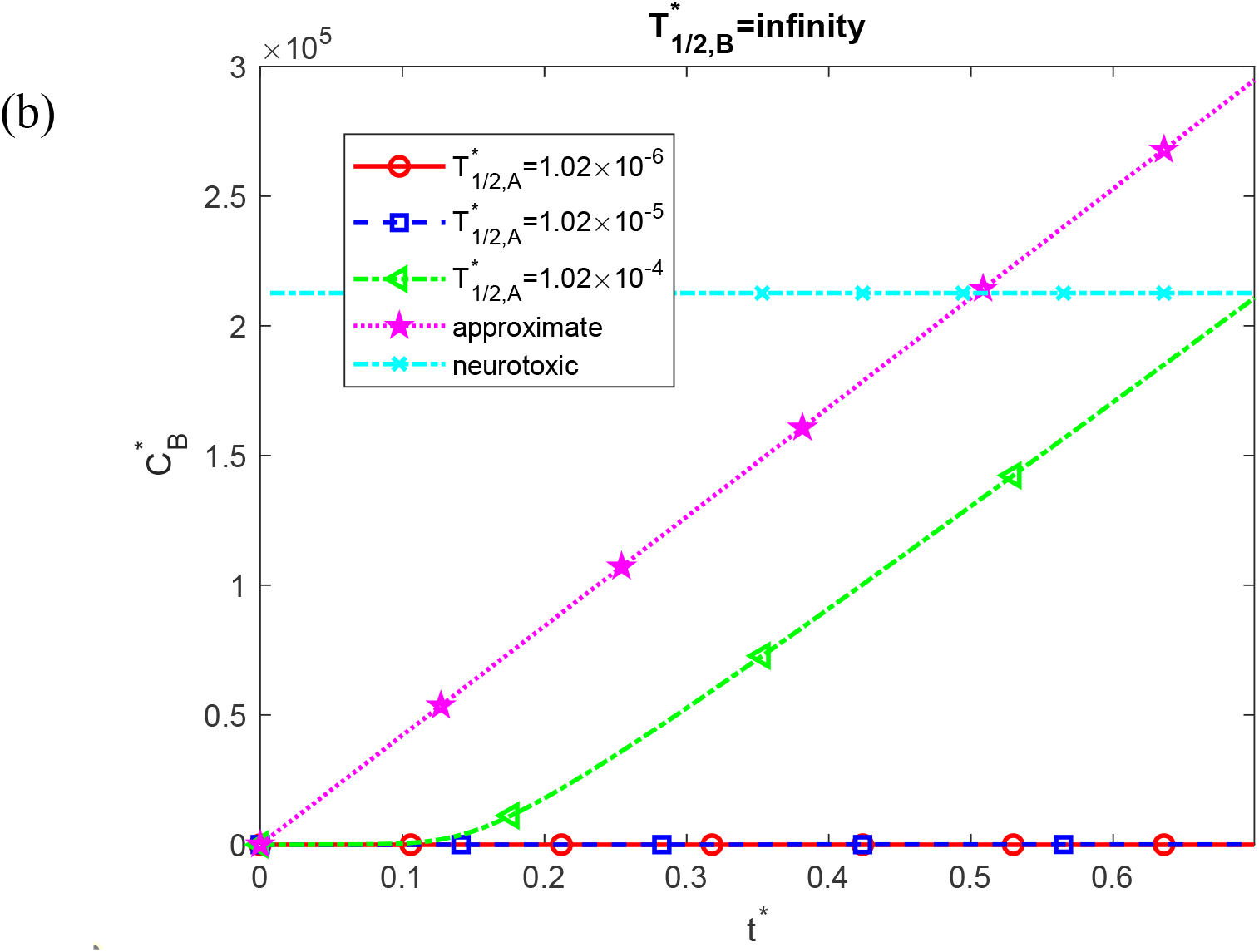
Dimensionless molar concentrations of Aβ monomers, 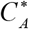 (a) and Aβ aggregates, 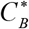 (b) as a function of dimensionless time, *t*^*^. Approximate solutions for 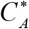 and 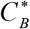 are computed using Eqs. (S11) and (S10), respectively. The approximate solution for 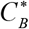 is directly proportional to *t*^*^. These results are presented at the right-hand side boundary of the CV, *x*^*^ =1, and correspond to the scenario with an infinite half-life of Aβ aggregates, 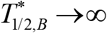. Results are shown for three values of the dimensionless half-life of Aβ monomers, 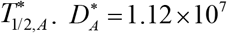 and 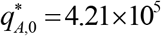.

**Fig. S9.**
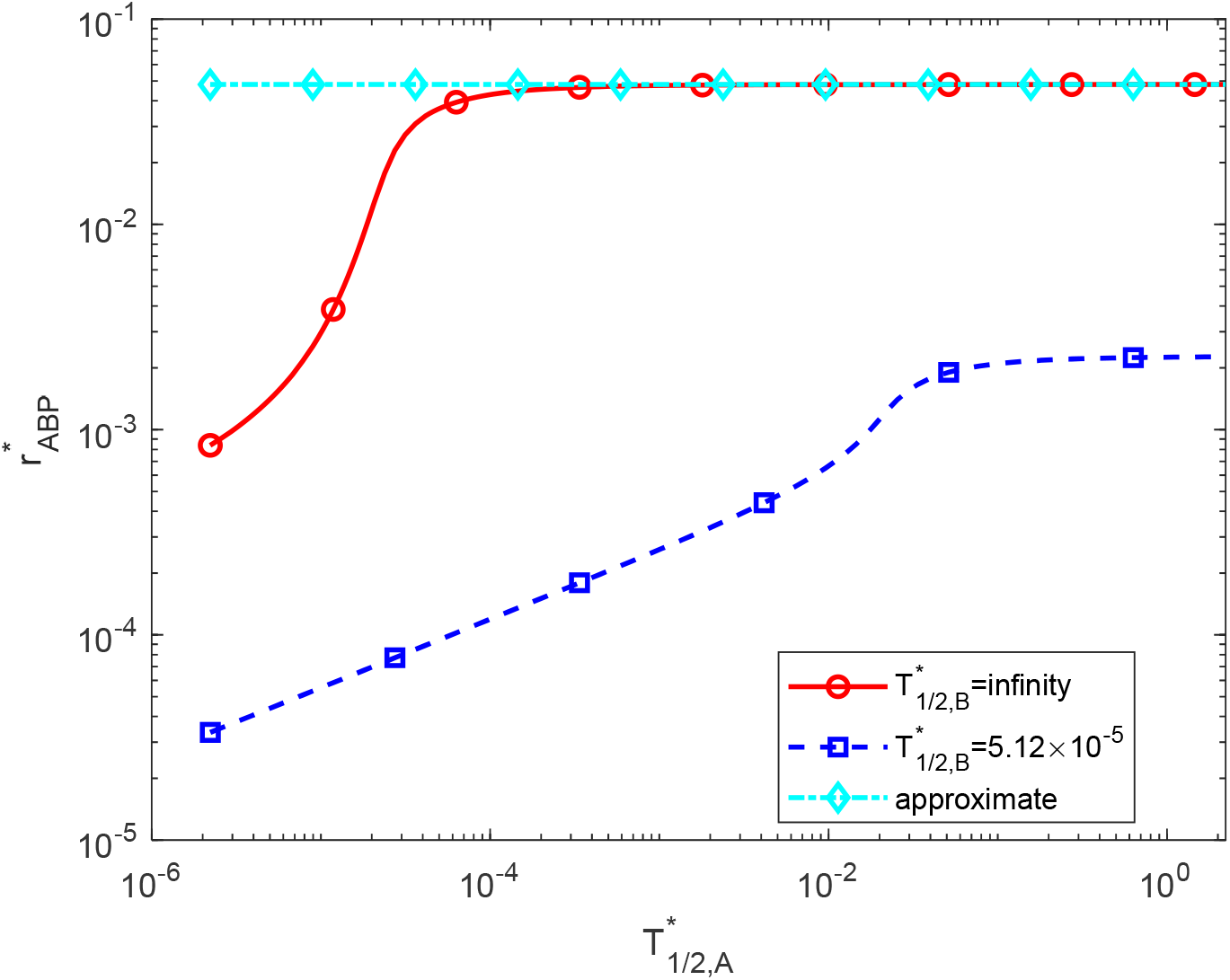
Dimensionless radius of an Aβ plaque, 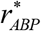, as a function of the dimensionless half-life of Aβ monomers, 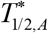, at 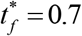 (10 years). Dimensional radius of the Aβ plaque corresponding to the largest value of 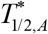 in Fig. S9 for infinite half-life of Aβ aggregates is 0.048 (corresponding dimensional radius is 2.40 μm). 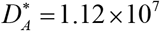 and 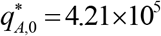.

**Fig. S10.**
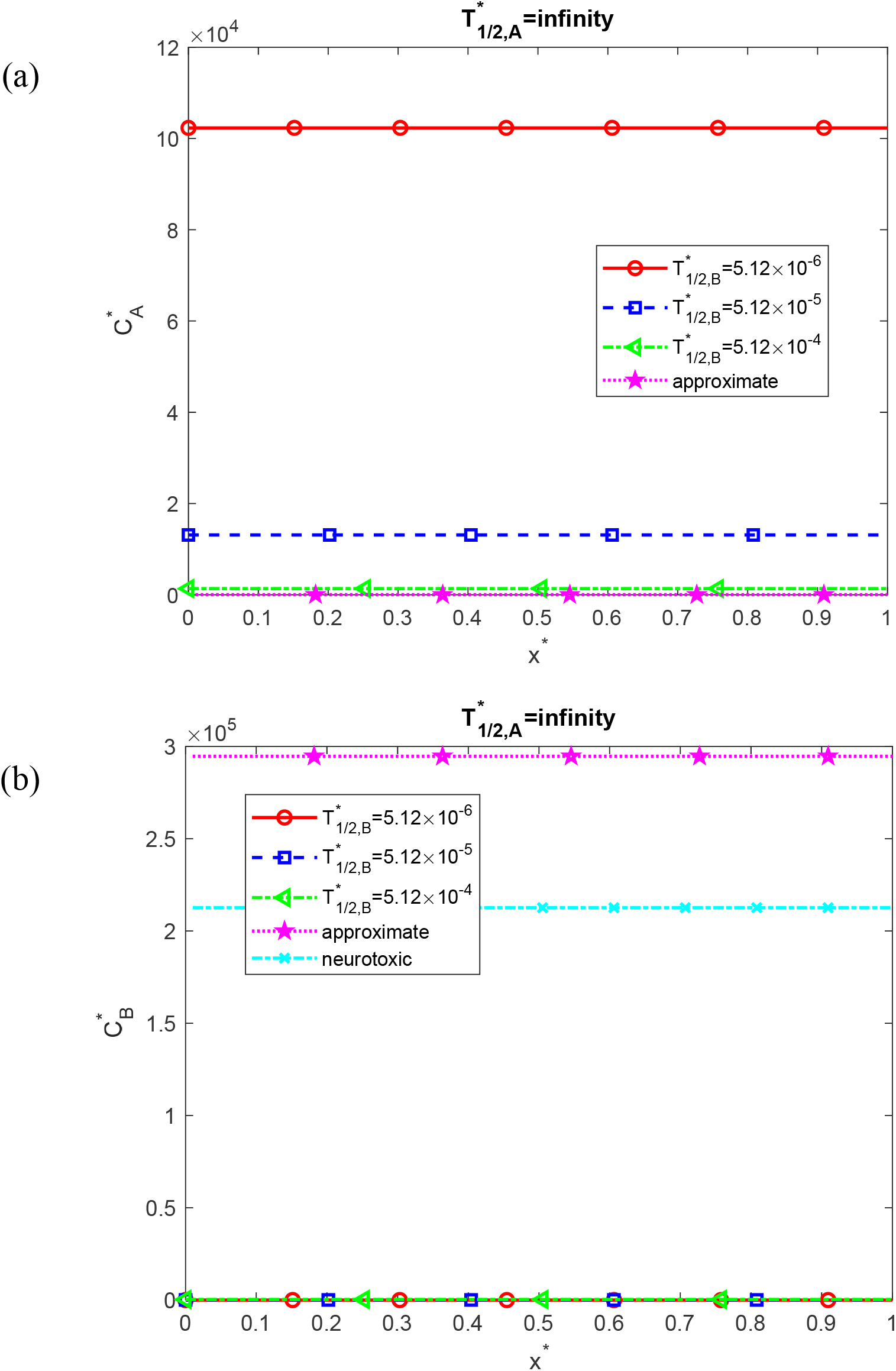
Dimensionless molar concentrations of Aβ monomers, 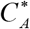 (a) and Aβ aggregates, 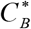 (b) as a function of the dimensionless distance from the surface releasing Aβ monomers, *x*^*^. Approximate solutions (independent of *x*^*^) for 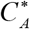 and 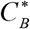 are computed using Eqs. (S11) and (S10), respectively. These results are presented at 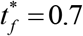 (10 years) and correspond to the scenario with an infinite half-life of Aβ monomers, 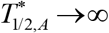. Results are shown for three values of the dimensionless half-life of Aβ aggregates, 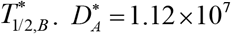 and 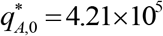.

**Fig. S11.**
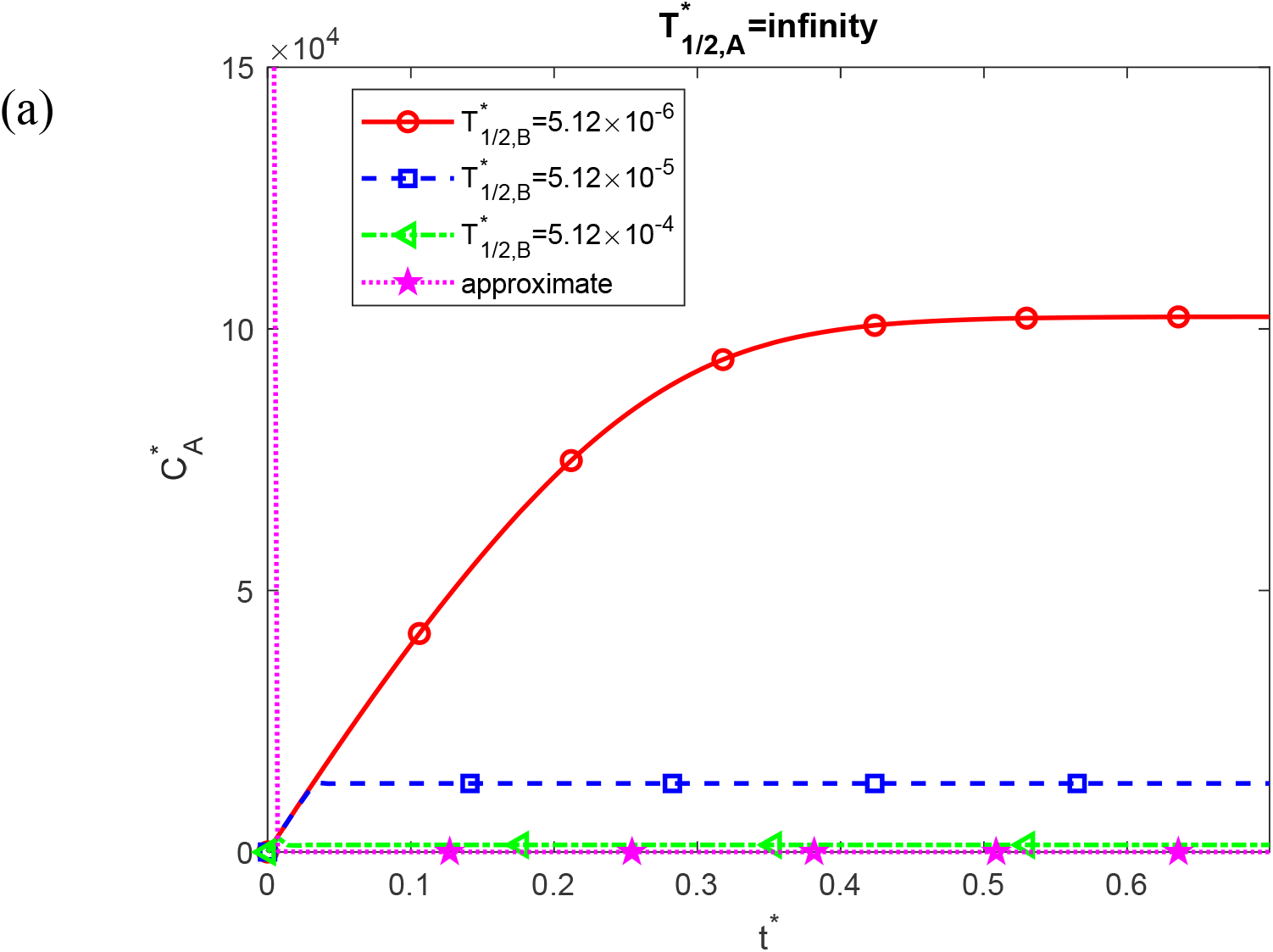
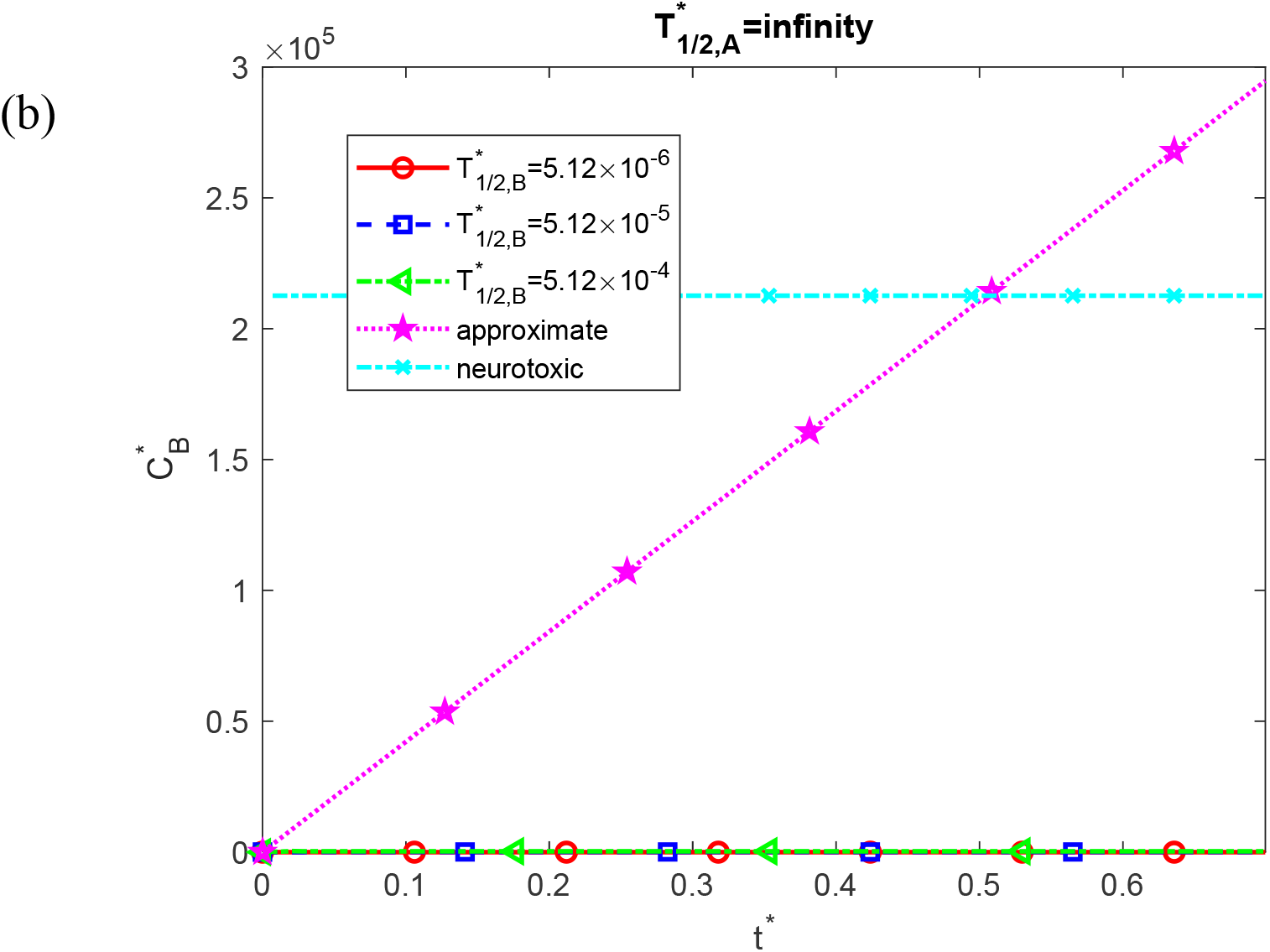
The dimensionless molar concentrations of Aβ monomers, 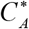 (a) and Aβ aggregates, 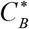 (b) as a function of dimensionless time, *t*^*^. The approximate solutions for 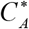 and 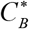 are determined using Eqs. (S11) and (S10), respectively. The approximate solution for 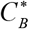 is directly proportional to *t*^*^. These computational results are presented at the right-hand side boundary of the CV, *x*^*^ =1, depicting the scenario with an infinite half-life of Aβ monomers, 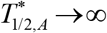. The results are presented for three different values of the dimensionless half-life of Aβ aggregates, 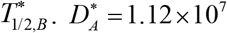 and 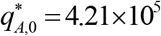.

**Fig. S12.**
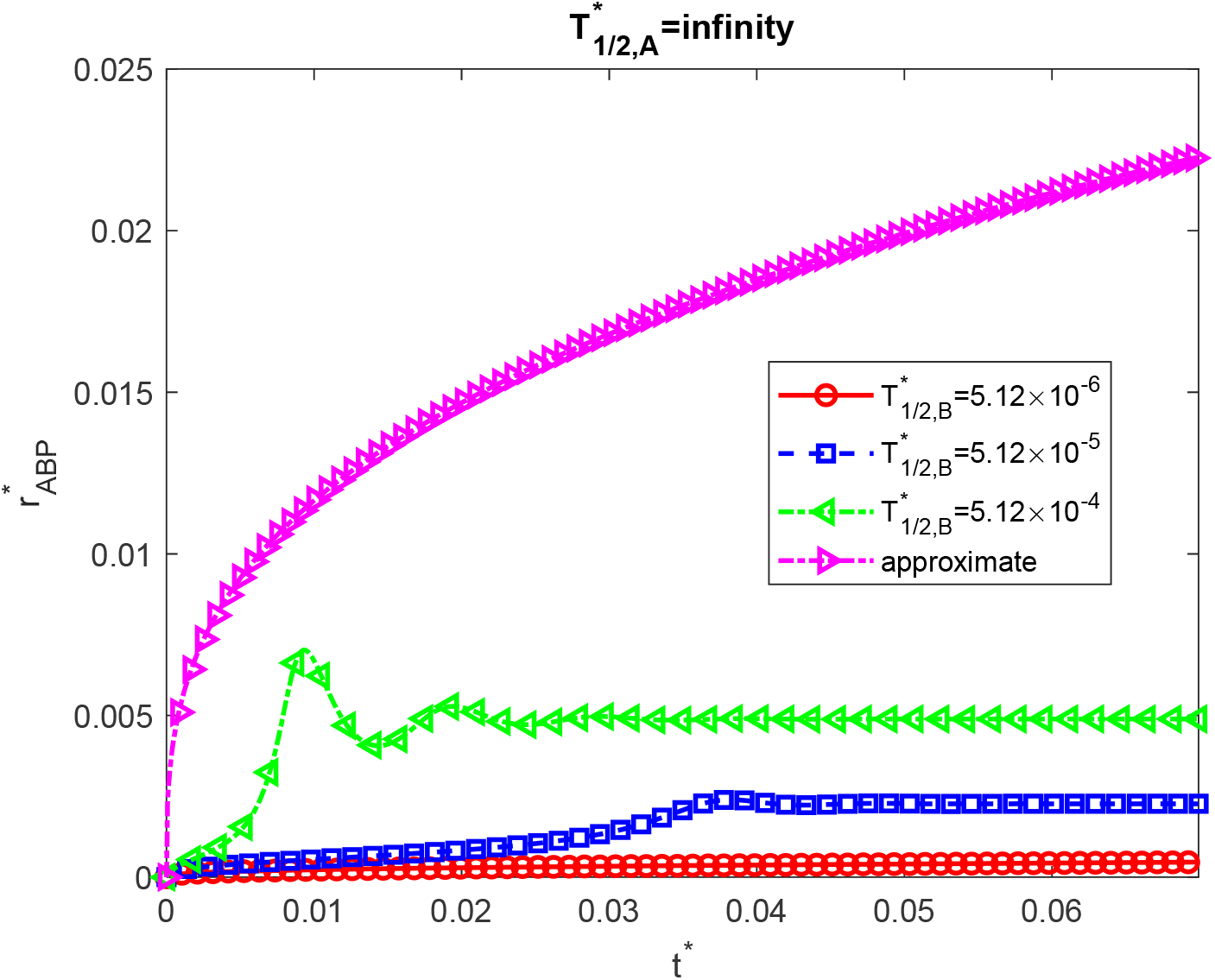
Similar to Fig. 9b, this figure is computed with a mesh ten times finer with respect to *t* ^*^ and *x*^*^ than Fig. 9b.

**Fig. S13.**
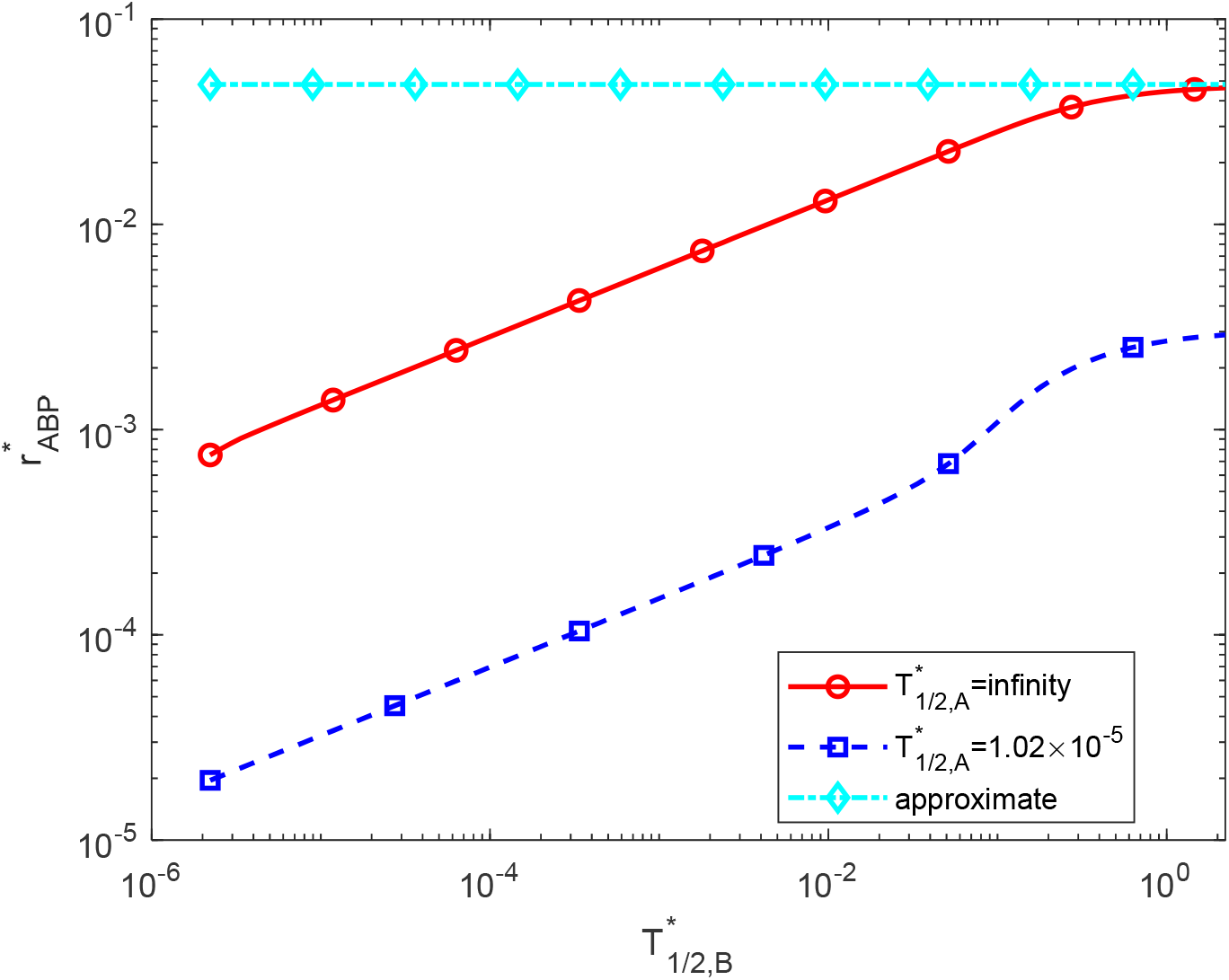
The dimensionless radius of a growing Aβ plaque, 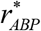, as a function of the dimensionless half-life of Aβ aggregates, 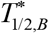, at 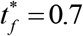 (10 years). The dimensionless radius of the Aβ plaque corresponding to the largest value of 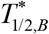 in Fig. S13 for the infinite half-life of Aβ aggregates is 0.046 (corresponding dimensional radius is 2.32 μm). 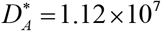 and 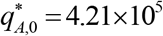.

